# Mechanism of fatty acid uptake and inhibition in human FATP2

**DOI:** 10.64898/2026.07.06.736331

**Authors:** Zhenglai Zhang, Mengqing Zhou, Yuwen Huang, Wencheng Wu, Haizhan Jiao, Minxian Dai, Taizhen Liang, Jiayue Wen, Ziqi Cheng, Xiancai Ma, Jiao Yuan, Hongli Hu, Jinsai Shang, Ronen Marmorstein, Xuepeng Wei

## Abstract

Fatty Acid Transport Protein 2 (FATP2) couples fatty acid uptake to intracellular activation and is associated with pathological lipid accumulation in cancer and nonalcoholic fatty liver disease. Here, we present cryo-electron microscopy structures of human FATP2 across its reaction cycle. Our structures suggest that FATP2 recruits fatty acids directly from the membrane interface through a hydrophobic tunnel. Catalysis involves a ∼130° rotation of the C-terminal domain, a transition trapped by the antihypertensive drugs isradipine and benidipine. Both drugs lock the enzyme in a thioester-forming state, but benidipine exhibits superior efficacy by extending a bulky moiety into the primary catalytic tunnel to sterically block substrate entry. Furthermore, we identify a product inhibition mechanism where excess acyl-CoA traps the enzyme, potentially limiting metabolic overload. These findings provide a structural framework for understanding vectorial fatty acid channeling and a scaffold for developing modulators of metabolic flux.

**Highlights:**

- Cryo-EM structures of human FATP2 reveal a membrane-anchored lollipop topology
- Endogenous fatty acids within a hydrophobic tunnel delineate the fatty acid uptake pathway
- Isradipine and benidipine displace fatty acids to trap a non-productive conformation
- Acyl-CoA product inhibition may provide negative feedback via steric occlusion

## Introduction

Long-chain fatty acids are vital for membrane biogenesis, energy storage, and signaling, ^1,2^ but their insolubility poses a fundamental challenge as they partition into membranes before conversion into metabolically committed intermediates. Nature has solved this problem by activating chemically inert free fatty acids to acyl-CoA thioesters to enter metabolic pathways.^3^ Dysregulation of this metabolic flux drives common human diseases such as fatty liver disease and cardiometabolic dysfunction.^4,5^ Lipid remodeling in the tumor microenvironment can also fuel immune suppression and tumor progression.^6–8^ Central to these processes is the Fatty Acid Transport Protein family, also known as FATPs or SLC27, particularly FATP2 (SLC27A2), which couples cellular uptake directly to activation through its intrinsic very long-chain acyl-CoA synthetase (VLACS) activity.^3,9–12^ This enzymatic activity is also clinically significant because liver-enriched FATP2 thioesterifies the prodrugs bempedoic acid and EVT0185 into their active CoA metabolites, bempedoyl-CoA and EVT0185-CoA, which are used to treat hypercholesterolemia by inhibiting ATP citrate lyase.^13–15^ While this highlights the enzyme’s pharmacological versatility, its primary physiological role is to regulate endogenous lipid flux. Because FATP2 directly controls the rate that membrane-derived fatty acids are committed to downstream pathways, its activity provides a direct point of physiological and pharmacological control. Consistent with this role, elevated or dysregulated FATP2 is strongly linked to pathological lipid accumulation and cancer-promoting lipid programs, making it a highly relevant clinical node for therapeutic intervention.^8,16^

Despite its therapeutic potential, targeting FATP2 is hampered by unresolved mechanistic questions at the core of its biology: how it couples membrane-associated fatty acid acquisition directly to ATP-dependent acyl-CoA production, and whether it exhibits the fatty acid transport activity implied by its solute carrier (SLC) designation. ^9,17^ The "vectorial acylation" model suggested that inhibiting FATP2 could simultaneously restrict substrate availability and limit the production of bioactive lipids. The SLC classification was historically supported by hydropathy analysis and experimental characterization of murine FATP1 and yeast FATP homologs, where epitope tagging suggested a complex transporter architecture with a polytopic membrane topology and extracellular N-terminus.^17–20^ While SLC proteins typically function via alternating-access mechanisms facilitated by multiple transmembrane domains, AlphaFold structural predictions depict FATP2 with only a single transmembrane domain, an architecture that contrasts sharply with the multi-pass transporters.^21^ Moreover, FATP2 contains an AMP-binding motif characteristic of adenylate-forming acyl-CoA synthetase enzymes, which catalyze a two-step reaction: the ATP-dependent adenylation of the carboxylate substrate followed by the formation of a thioester with Coenzyme A.^10,22^ Consequently, distinguishing between this enzyme-driven uptake model, where ATP drives thermodynamic capture, and an intrinsic transport model is crucial. In this latter scenario, the protein would utilize specific conformational changes to physically shuttle fatty acids across the bilayer independent of its enzymatic activity. Regarding its unusual enzymatic activity, how FATP2 facilitates the uptake of fatty acids from the membrane environment to form long-chain acyl-CoAs remains unresolved. Therefore, resolving whether FATP2 operates via a physical translocation mechanism or a purely enzymatic strategy is an important mechanistic question in the field and is also a vital prerequisite for rational drug design.

To resolve this, we present single-particle cryo-electron microscopy structures of full-length human FATP2 captured across its reaction cycle, including complexes with the pharmacological substrate bempedoic acid, the product palmitoyl-CoA, and the calcium channel blockers isradipine and benidipine,^23,24^ which we identify herein as novel FATP2 inhibitors. Our structures do not support a multi-pass, alternating-access transporter architecture, instead revealing a lollipop-like shape anchored by a single transmembrane tether and a membrane-associated amphipathic helix (AH). Crucially, this architecture features a continuous hydrophobic tunnel connecting the membrane interface directly to the cytosolic catalytic center, facilitating a "vectorial channeling" mechanism of direct substrate extraction and conversion to acyl-CoA. Mechanistically, we identify distinct conformational states defined by a ∼130-degree rotation of a flexibly tethered C-terminal domain required to transition from the initial adenylate-forming conformation to the subsequent thioester-forming state, alongside an unexpected product feedback mechanism where acyl-CoA occupies the ATP-binding site and occludes the fatty acid tunnel to trap the protein in a non-productive state. Finally, we characterize the structural basis of inhibition, revealing that these drugs act as potent inhibitors by displacing a membrane-interface fatty acid while simultaneously trapping the enzyme in a distinct, non-productive thioester-forming conformation. While sharing this binding mode, benidipine uniquely bears an extended substituent that penetrates the primary catalytic tunnel, imposing severe steric occlusion. These findings establish a structural framework for how FATP2 couples fatty acid uptake to CoA activation, providing a solid foundation for drug discovery to modulate fatty acid flux in disease.

## Results

### Substrate specificity and structure of human FATP2

To investigate the substrate preference of FATP2 for fatty acid chain length (Figure 1B), we challenged the purified full-length protein from HEK293F cells (Figure S1) with a diverse library of saturated and unsaturated fatty acids, as well as the pharmacological agent bempedoic acid. The enzyme displayed a striking specificity profile: while largely inert toward short-chain species such as butyrate (C4:0), it exhibited robust activity across a range of medium-, long-, and very-long-chain fatty acids (Figure 1B). Significant activity was observed with saturated species like palmitate (C16:0) and stearate (C18:0), as well as unsaturated forms including palmitoleate (C16:1), oleate (C18:1), erucate (C22:1), and nervonate (C24:1). Notably, *in vitro* activity peaked with decanoate (C10:0), surpassing rates observed for physiological long-chain substrates. Furthermore, FATP2 demonstrated high enzymatic activity toward bempedoic acid (BEM), a 19-carbon dicarboxylic acid structurally analogous to long-chain fatty acids, comparable to its maximal activity against natural fatty acid substrates (Figure 1B).

**Figure 1.**
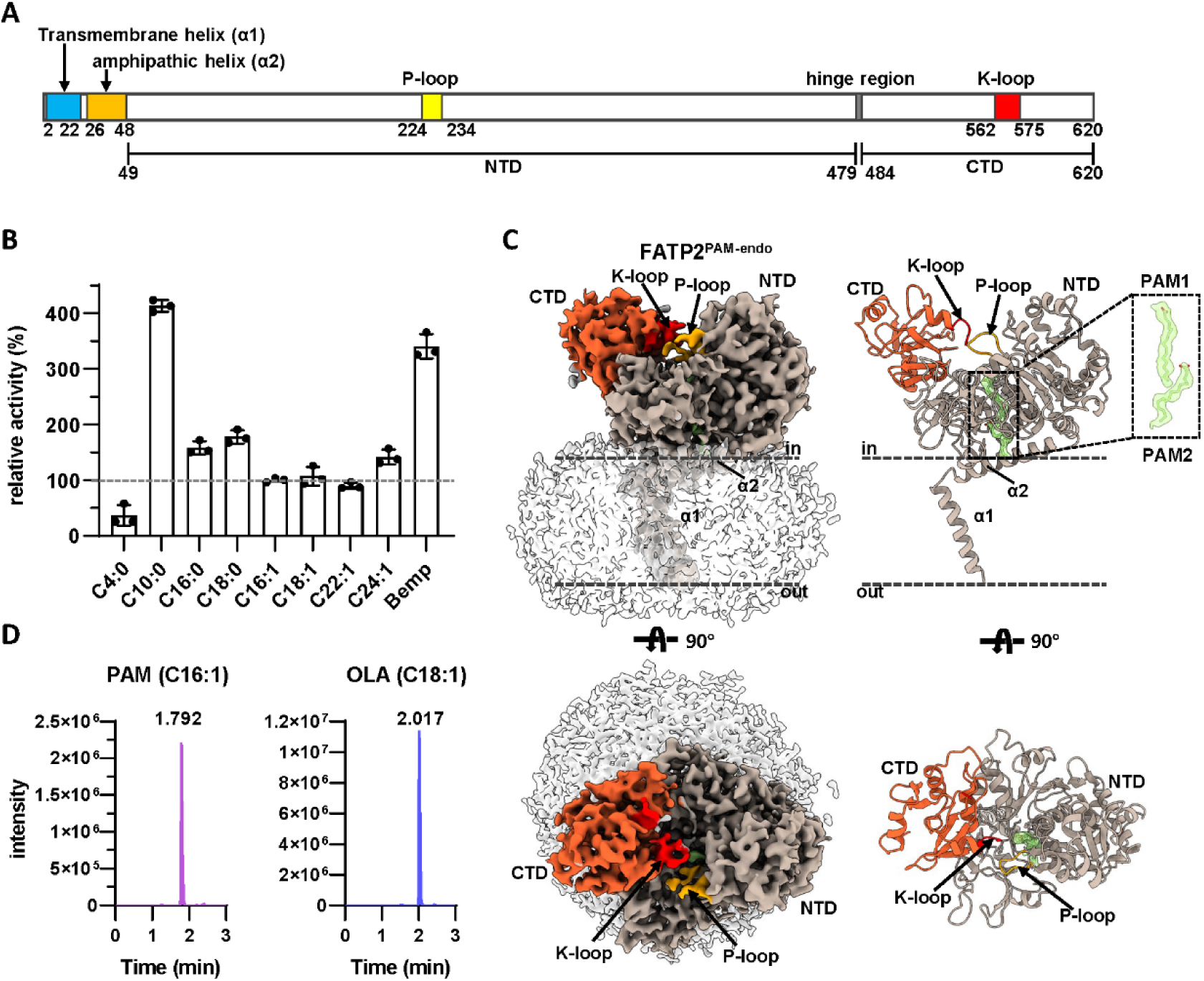
Functional characterization and structural organization of human FATP2. (A) Domain organization of human FATP2. Key structural elements include the N-terminal transmembrane helix (α1; blue), the amphipathic helix (α2; orange), the ATP-binding P-loop (yellow), the flexible hinge region (grey), and the lysine-rich K-loop (red). (B) Substrate specificity profile of FATP2. Relative acyl-CoA synthetase activity was measured against fatty acids of varying chain lengths (C4:0–C24:1) and the prodrug bempedoic acid (BEM). Data represent mean ± s.d. (normalized to C16:1 fatty acid, n ≥3 technical replicates). (C) Cryo-EM structure of the FATP2–endogenous fatty acid complex (FATP2^PAM-endo^). Left: Surface representation of the cryo-EM density map (side view top, top view bottom), with the detergent micelle shown as a grey transparent surface. Right: Cartoon representation of the atomic model, colored to distinguish the N-terminal domain (NTD, grey), C-terminal domain (CTD, orange), P-loop (yellow), and K-loop (red). The inset displays the cryo-EM density (green transparent surface) corresponding to two bound fatty acids PAM1 and PAM2, with PAM1 positioned inside the hydrophobic tunnel and PAM2 located close to the membrane. Dashed lines indicate the approximate boundaries of the membrane. (D) Identification of endogenous ligands. LC–MS analysis of lipids extracted from the purified protein reveals the enrichment of C16:1 and C18:1 fatty acids. See also Figures S1-S6.

To elucidate the structural mechanism enabling the accommodation of such variable chain lengths and bulky analogs, we determined the cryo-EM structure of FATP2 at 3.2 Å resolution (Figures 1C; Figures S2 and S3). The resolved architecture reveals a lollipop-like assembly comprising a large cytosolic catalytic body and a membrane-anchoring tether (Figures 1A and 1C). The cytosolic region is segmented into a large N-terminal domain (NTD; residues 49–479) and a smaller, flexible C-terminal domain (CTD; residues 484–620), connected by a short hinge (residues 480–483), characteristic of acyl-CoA synthetase enzymes.^25^ Within this cytosolic architecture, the NTD harbors the conserved P-loop (phosphate-binding loop; residues 224-234) essential for ATP coordination. However, the distinct K-loop (lysine-rich loop, residues 562-575) exhibits weak density in this EM map, likely due to its inherent flexibility in the absence of ATP binding. The protein is secured to the membrane by a single N-terminal transmembrane helix (α1; residues 2–22) followed by an amphipathic helix (α2, residues 26-48). The cryo-EM density reveals a detergent micelle belt surrounding this region, delineating the likely position of the lipid bilayer where α2 adopts a tilted orientation at the interface (Figure 1C). This topology stands in sharp contrast to canonical Solute Carrier (SLC) transporters, which typically feature a core of multiple transmembrane helices that form a central pore to facilitate substrate translocation via alternating-access mechanisms.^26–28^ Indeed, a structural comparison with the prototypical glucose transporter GLUT1 (SLC2A1) ^29^ and the representative lipid transporter MFSD2A ^30^ highlights the absence of such a transmembrane pore in FATP2, strongly implying that it does not function as a canonical transporter (Figure S4).

Inspection of the map revealed a substantial hydrophobic tunnel traversing the NTD, occupied by two separate continuous densities resembling fatty acids (Figure 1C). Lipidomic analysis of the purified complex identified the presence of palmitoleic acid (C16:1, PAM) and oleic acid (C18:1, OLA) (Figure 1D), corresponding to a 37- and 29-fold enrichment, respectively, relative to the buffer control (Figure S5). Given the similar chain lengths of these predominant species and the resolution limits, we modeled both ligands as PAM. We assigned the prominent density in the upper tunnel facing the cytosol as PAM1 and the secondary density near the amphipathic helix at the putative membrane interface as PAM2 (Figure 1C). Thus, the protein was captured in a fatty acid-bound state, hereafter referred to as FATP2^PAM-endo^.

A structural homology search within the PDB identified the plant 4-coumarate:CoA ligase (4CL2) as a close structural homolog of FATP2 belonging to the acyl-CoA synthetase family. We therefore superimposed our FATP2^PAM-endo^ structure onto the structures of the plant 4CL2 in both adenylate-forming (PDB code: 5BSM) and thioester-forming (PDB code: 5BSR) states.^31^ This comparison yielded RMSD values of 2.66 Å and 2.75 Å, respectively (Figure S6), confirming a conserved global fold. However, closer inspection of the C-terminal domain (CTD) orientation reveals that FATP2^PAM-endo^ specifically adopts a nucleotide-free adenylate-forming conformation. This suggests that FATP2 is inherently primed for catalysis by effectively pre-organizing the machinery for the initial half-reaction.

Consistent with its physiological function, the FATP2 tunnel is substantially deeper than that of medium-chain acyl-CoA synthetase ACSM2A,^32^ which processes medium-chain fatty acids, providing the structural basis of long-chain substrate processing (Figure S7). Importantly, the two fatty acid densities, PAM2 near the membrane interface (site 2) and PAM1 deep within the upper tunnel (site 1), suggest a directional path for fatty acid movement (Figure 1C). We propose that fatty acids partition into the membrane, equilibrate into the inner leaflet, enter the tunnel near α2 at the cytosolic membrane interface, and are then guided to the active site (site 1) for activation.

The apparent preference for decanoate (C10) in vitro likely reflects substrate availability and partitioning in our assay conditions rather than physiological specialization. Short-chain substrates such as C4 may be insufficiently hydrophobic to displace endogenous lipids or to occupy the deep tunnel stably. Conversely, very long-chain fatty acids can aggregate in aqueous buffer, reducing the effective concentration accessible to the enzyme. Thus, C10 likely represents a practical optimum: hydrophobic enough to enter the tunnel but soluble enough to remain available.

### Recognition of ATP and prodrug bempedoic acid

To provide a structural basis for the ATP- and acyl-substrate dependent adenylation half-reaction and the recognition of pharmacologic substrates, we determined the cryo-EM structure of FATP2 in the presence of excess ATP and the prodrug bempedoic acid (BEM) at 2.6 Å resolution (hereafter FATP2^ATP-BEM^; Figures 2A-2B; Figure S3). Brief incubation of the complex on ice minimized enzymatic turnover, allowing us to capture the pre-catalytic adenylate-forming conformation. This reveals that the fatty acid substrate resides within the hydrophobic tunnel, mirroring the binding mode of endogenous fatty acid. Directly adjacent to this tunnel, the ATP cofactor binds at the interface of the NTD and CTD, an arrangement that positions the reactants in proximity for catalysis (Figures 2A-2B). A structural superposition of this catalytic complex with the nucleotide-free adenylate-forming conformation (FATP2^PAM-endo^) reveals a high degree of overall similarity with an RMSD of 0.93 Å.

**Figure 2.**
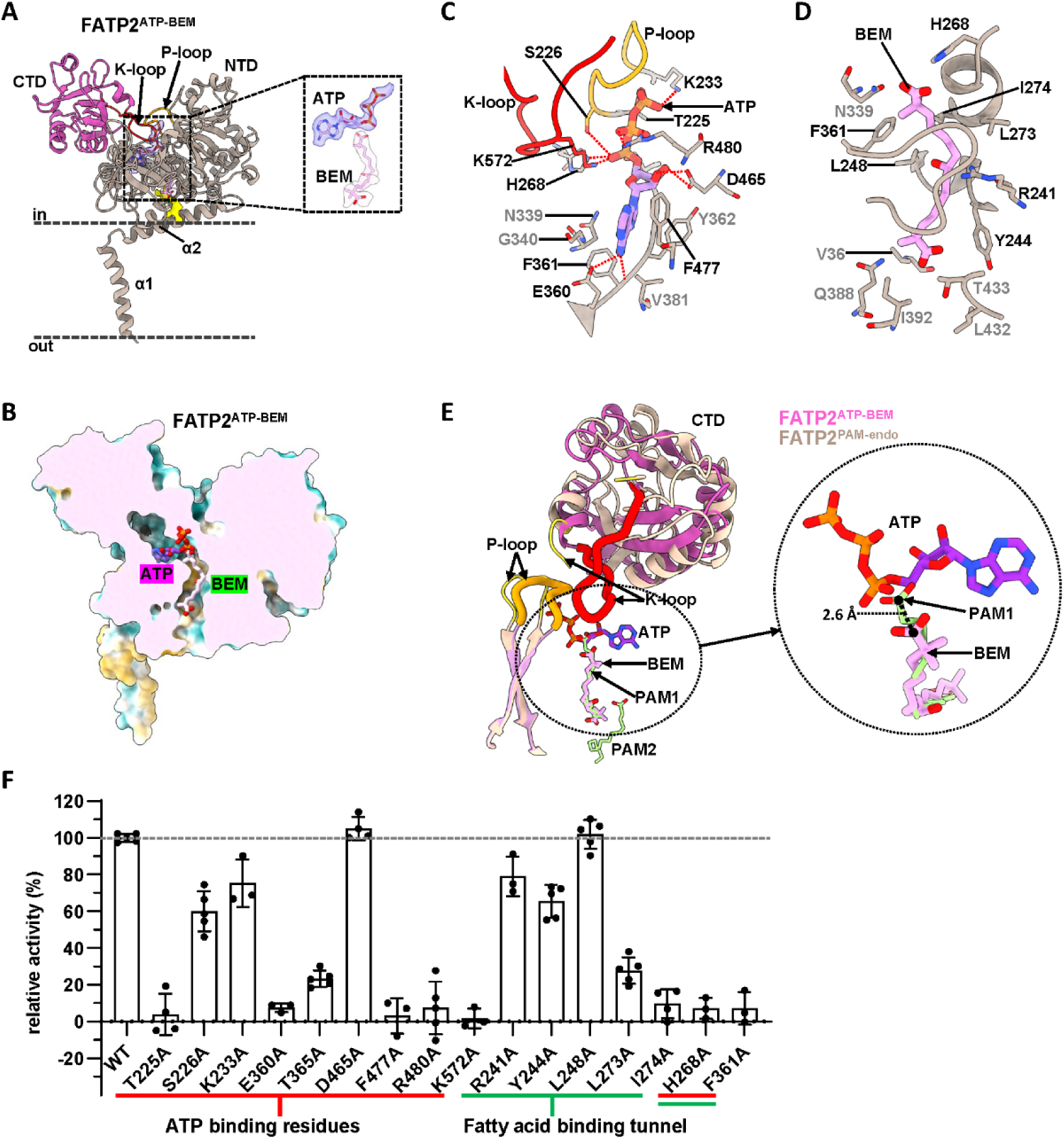
Structural basis of ATP and bempedoic acid recognition by FATP2. (A) Overall architecture of the FATP2^ATP–BEM^ complex. The N-terminal domain (NTD) is colored grey, and the C-terminal domain (CTD) is pink. The P-loop and K-loop are highlighted in yellow and red, respectively. The inset displays the cryo-EM density map for ATP and bempedoic acid (BEM). (B) Cut-away surface representation revealing the ATP-binding pocket and the hydrophobic substrate tunnel occupied by BEM. (C) Detailed view of the ATP-binding pocket. Key residues involved in nucleotide recognition are shown as sticks, with hydrogen bonds and salt bridges indicated by red dashed lines. (D) Detailed view of the substrate-binding tunnel accommodating bempedoic acid. Residues lining the hydrophobic channel are shown as sticks. (E) Structural superposition of the FATP2^ATP–BEM^ complex (magenta/pink) and the nucleotide-free FATP2^PAM-endo^ complex (tan). The magnified view highlights the 2.6 Å displacement of the substrate carboxyl group (from the position of PAM1 to BEM) induced by ATP binding. (F) Mutational analysis of key residues within the ATP-binding site and substrate tunnel. Enzymatic activity was measured for wild-type (WT) and mutant FATP2. Underlines categorize residues by structural role: ATP recognition (red), tunnel composition (green), or both (double lines). Data are presented as mean ± s.d. (n ≥3 technical replicates). See also Figures S3 and S8.

Despite this conserved global architecture, distinct local conformational changes are observed within the substrate-binding site. Bempedoic acid occupies the substrate-binding tunnel, effectively replacing the primary palmitoleic acid (PAM1) observed in the nucleotide-free state. This confirms that FATP2 recognizes bempedoic acid as a substrate for CoA activation within this hydrophobic tunnel, which extends from the active site to the membrane-facing surface (Figure 2B). While a second palmitoleic acid density (PAM2) was observed in the nucleotide-free structure, it is poorly resolved in the FATP2^ATP-BEM^ complex, likely due to low occupancy or high flexibility in the presence of the prodrug and nucleotide. Notably, the bempedoic acid molecule is shifted approximately 2.6 Å relative to the position of PAM1 in the nucleotide-free structure (Figure 2E). This deviation appears to be driven by the bulky ATP molecule, which binds at the top of the active site and acts as a steric wedge, forcing the reactive carboxyl group of bempedoic acid to shift to a lower, catalytically competent position relative to the PAM1 observed in the FATP2^PAM-endo^ structure.

Structural and mutational analyses delineate a coordinated network within the FATP2 active site, revealing distinct roles for nucleotide coordination and substrate positioning (Figures 2C-2D, and 2F). ATP binding is orchestrated by the N-domain P-loop (residues 224–234) and the C-domain K-loop (residues 562–575), which enclose the triphosphate moiety (Figure 2C). Consistent with this model, disrupting the P-loop (T225A) or the K-loop (K572A) abolished catalytic activity, reflecting the role of the K-loop in sealing the active site and neutralizing the α-phosphate charge (Figure 2F). In contrast, the peripheral P-loop variants S226A and K233A retained substantial function. Similarly, the strict requirement for adenine and ribose stabilization was confirmed when alanine substitutions of E360, F477, and R480 resulted in a near-complete loss of activity, while T365A exhibited significantly reduced activity; conversely, the D465A mutant remained active, suggesting the hydrogen bond between D465 and the ribose is functionally redundant.

Bridging the nucleotide and fatty acid binding regions, residues F361 and H268 exhibit critical bifunctional roles. The backbone of F361 secures the adenine base, while its hydrophobic side chain forms an integral part of the fatty acid binding tunnel. Accordingly, the F361A mutation resulted in a near-complete loss of enzyme function. H268 is equally pivotal as it positions the reactive carboxyl group of bempedoic acid for adenylation while simultaneously stabilizing the ATP α-phosphate through a hydrogen bond. This dual coordination is essential for catalysis, as evidenced by the similarly impaired activity of the H268A mutant. These functionally critical residues are conserved across the ANL enzyme family, consistent with a shared catalytic architecture (Figure S8). The fatty acid binding tunnel features a hydrophobic interior and electropositive regions. Mutations to tunnel-lining residues (L273A, I274A) significantly reduced activity, whereas variants R241A and Y244A caused only moderate reductions.

### Structural basis of product inhibition

To investigate the structural basis of product formation and release, we determined the 2.7 Å cryo-EM structure of FATP2 incubated with its reaction products, AMP and palmitoyl-CoA (FATP2^Product^). Although AMP was included during incubation, only palmitoyl-CoA is resolved in the cryo-EM density (Figures 3A-3B), whereas AMP is not resolved. Additionally, we determined the structure of FATP2 bound to Coenzyme A alone (FATP2^CoA^) at 3.3 Å resolution to serve as a reference for the second half-reaction’s substrate-bound state (Figures 3C-3D; Figure S2). Notably, despite their distinct incubation conditions, both the product-bound and CoA-bound structures adopt a conformation resembling the adenylate-forming state rather than the thioester-forming conformation required for the second half reaction.

**Figure 3.**
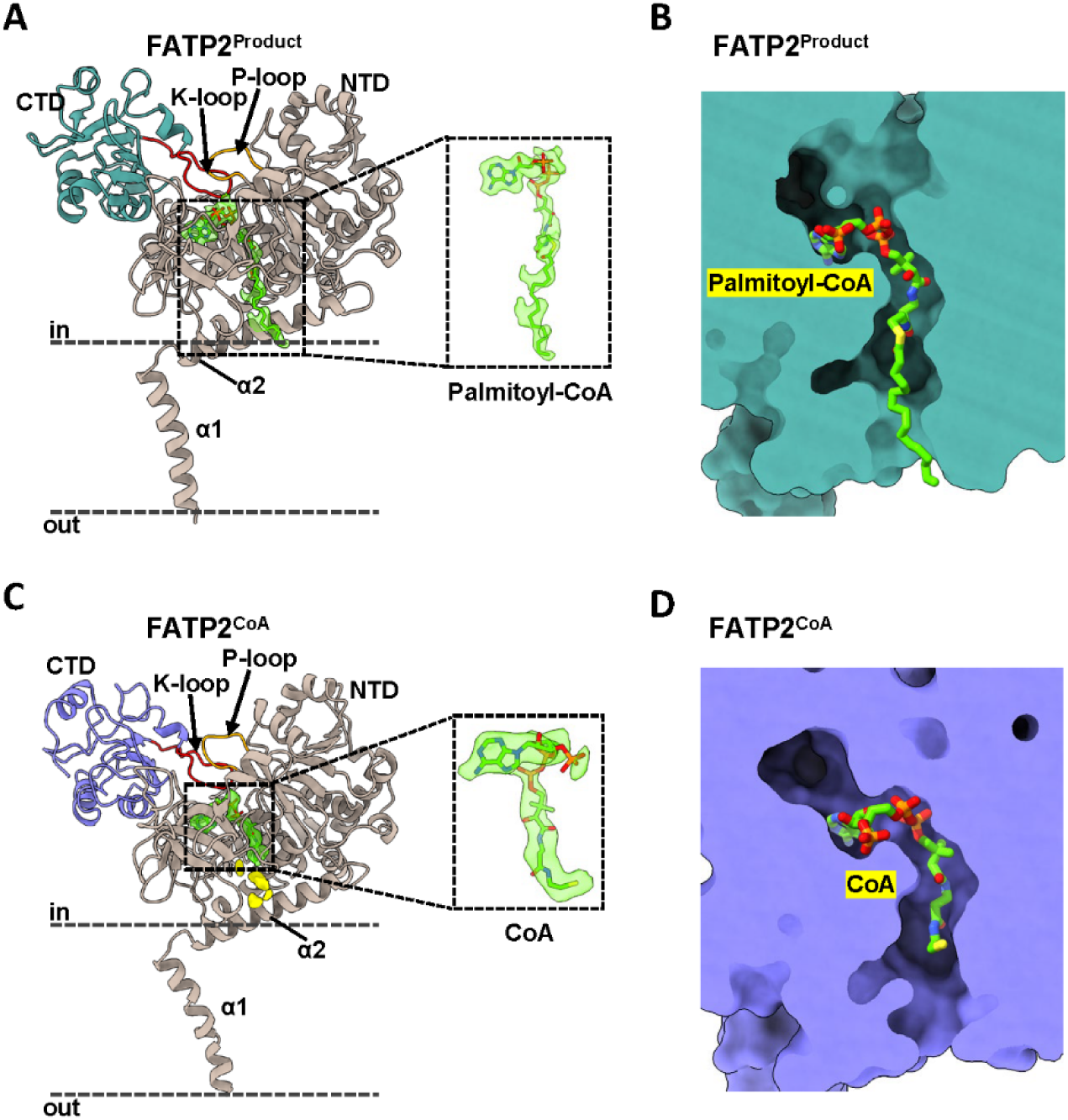
Structural basis of feedback inhibition in human FATP2. **(A)** Overall structure of the FATP2^Product^ complex. The P-loop and K-loop are colored orange and red, respectively. The dashed lines indicate the membrane boundaries. The inset displays the cryo-EM density of the palmitoyl-CoA molecule. **(B)** Cut-away surface representation of FATP2^Product^ revealing the palmitoyl-CoA binding tunnel. **(C)** Overall structure of the FATP2^CoA^ complex. The P-loop and K-loop are highlighted as in (A). The inset displays the cryo-EM density for the CoA molecule. **(D)** Cut-away surface representation of FATP2^CoA^ structure showing the CoA binding tunnel. See also Figure S9.

These structures reveal an extensive self-inhibition mode where the ligand essentially mimics the substrates but binds in a non-productive orientation that effectively jams the catalytic machinery. Specifically, the adenosine moiety of the CoA or palmitoyl-CoA molecule inserts itself directly into the nucleotide-binding pocket, occupying the precise site normally reserved for ATP (Figure 3). Concomitantly, the pantetheine arm extends into the fatty acid tunnel, and in the FATP2^Product^ complex the palmitoyl acyl chain further penetrates the tunnel. This dual-occupancy of CoA or Palmitoyl-CoA creates a severe steric blockade: the adenosine headgroup prevents ATP entry, while the tail portion physically obstructs the path for incoming fatty acids. Unlike a standard competitive inhibitor that might block just one site, this non-productive binding sterically occludes both the nucleotide binding pocket and fatty acid tunnel simultaneously.

These structural findings suggest that excess acyl-CoA can hijack the ATP and fatty acid binding sites, locking the enzyme in an inactive state to regulate metabolic flux. This model of steric occlusion is supported by structural studies of non-productive binding in human ACSM2A (Figure S9A) and aligns with kinetic evidence of substrate inhibition in *Leishmania* LiAcs1.^32,33^ Validating this feedback mechanism, we determined the IC_50_ of palmitoyl-CoA for FATP2 to be 60.1 µM (Figure S9B). This is also consistent with murine FATP1 studies, which showed progressive inhibition ranging from ∼20% at 10 µM to over 60% at 100 µM palmitoyl-CoA.^34^

### Isradipine and Benidipine trap FATP2 in the thioester-forming conformation

To identify therapeutic agents capable of targeting FATP2 with greater efficacy than the natural feedback mechanism, we screened a library of FDA-approved compounds (Figure S10). This screen identified two dihydropyridine calcium channel blockers, isradipine and benidipine, as significant hits. Both drugs potently inhibited FATP2 activity, with isradipine and benidipine displaying IC_50_ values of 7.9 μM and 3.1 μM, respectively (Figure 4A). This represents a substantial increase in potency compared to the endogenous feedback inhibitor, palmitoyl-CoA (IC_50_ of 60.1 μM; Figure S9B). Although these drugs are traditionally prescribed for hypertension to manage cardiovascular risk via calcium channel blockade,^24,35–37^ our findings reveal a distinct, high-affinity mechanism of action against FATP2. This superior potency relative to endogenous regulation supports the potential repurposing of these agents for FATP2-driven malignancies and metabolic dysregulation.

**Figure 4.**
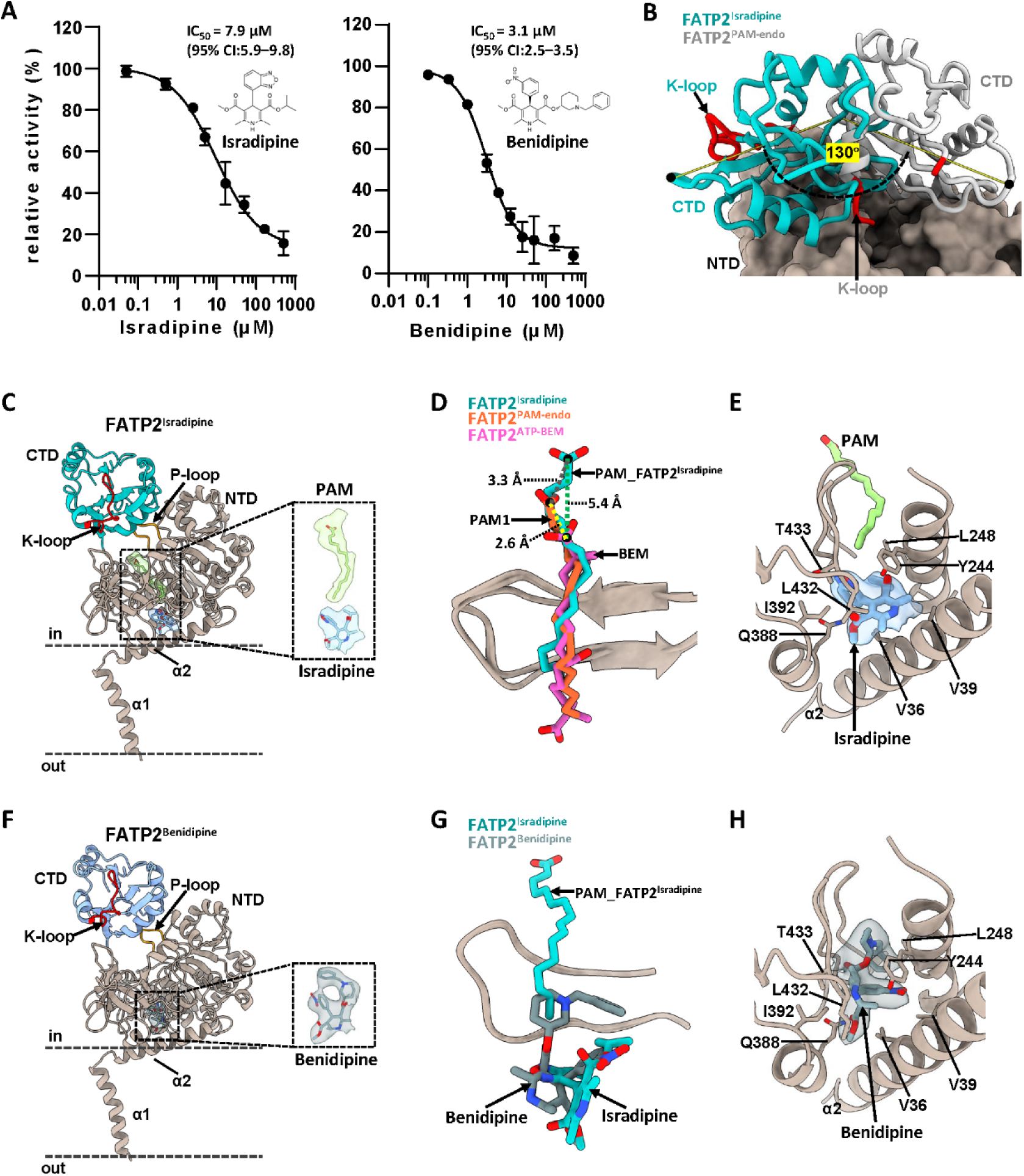
Identification and structural basis of isradipine and benidipine as FATP2 inhibitors. (A) Dose-response curve of isradipine and benidipine inhibition against FATP2, showing an IC_50_ of 7.9 μM and 3.1 μM, respectively. (B) Structural comparison of FATP2^Isradipine^ with FATP2^PAM-endo^ revealed a dramatic rotation of the C-domain of ∼130° relative to the N-domain. (C and F) Structure of FATP2 bound to isradipine (C) and benidipine (F) with ligand density highlighted. (D and G) Structural overlays of FATP2^Isradipine^, FATP2^PAM–endo^, and FATP2^ATP–BEM^ illustrating ligand displacement around the site 1 (D), and superposition of isradipine and benidipine bound to FATP2 (G). (E and H) Close-up view of the ligand-binding environment, showing the positions of PAM1 and isradipine in the FATP2^Isradipine^ complex (E) and the FATP2^Benidipine^ complex (H). Video S1. Conformational transition between adenylate-forming and thioester-forming conformations. Structural morph generated between the FATP2^PAM-endo^ and FATP2^Isradipine^ complexes, illustrating the ∼130° rotation of the C-terminal domain (CTD, cyan) relative to the stationary N-terminal domain (NTD, wheat). See also Figure S10.

To elucidate the molecular basis of dihydropyridine inhibition, we determined the cryo-EM structure of the isradipine-bound complex. Unlike prior datasets in this study that exclusively captured the adenylate-forming state, the cryo-EM data of FATP2^Isradipine^ resolved into two distinct populations (Figures S2, S10D and S10E). The first class resembled the adenylate-forming conformation; however, the map resolution was limited to 4.6 Å, likely due to drug-induced structural instability or heterogeneity (Figure S10D). Consequently, no model was built for this state. The second class, resolved at 3.5 Å resolution, captured the thioester-forming conformation and is hereafter referred to as FATP2^Isradipine^ (Figure S10E). Superposition of the FATP2^Isradipine^structure with FATP2^PAM-endo^ reveals a large-scale rearrangement: the C-terminal domain undergoes a rotation of approximately 130° relative to the N-terminal domain (Figure 4B; Video S1). This domain alternation is consistent with the mechanism reported for the related enzymes plant 4-coumarate:CoA ligase and human medium-chain acyl-CoA synthetase ACSM2A.^31,32^

Structural comparison reveals that isradipine occupies the secondary fatty acid site (site 2) observed in the FATP2–PAM-endo structure, as confirmed by the distinct ligand densities in this region (Figures 4C and 4E; Figure S11). Concomitantly, the primary fatty acid (PAM1) undergoes a marked positional shift within the catalytic tunnel. Whereas ATP binding in the active FATP2^ATP–BEM^ complex drives the ligand approximately 2.6 Å into an adenylation-competent pose (Figure 2E; Figure S11), the density for PAM1 in the isradipine-bound structure is shifted about 3.3 Å in the opposite direction away from this adenylation geometry (Figure 4D; Figure S11). This results in a total positional change of the fatty acid substrate PAM1 by 5.4 Å relative to its catalytically competent position (Figure 4D). Such a shift appears spatially incompatible with the compact interface of the adenylate-forming state. Consequently, we suggest that this steric alteration, evidenced by the variation in ligand densities across the endogenous, adenylate-forming, and inhibitor-bound states (Figure S11), may destabilize the resting conformation and bias the enzyme toward the non-productive thioester-forming state.

Given that both isradipine and benidipine inhibit FATP2, we reasoned that structurally related dihydropyridine analogues might exhibit similar effects and therefore performed a side-by-side comparison of isradipine with amlodipine and nifedipine. In contrast to isradipine, neither amlodipine nor nifedipine showed significant inhibition of FATP2 activity (Figure S10C). Although all four compounds share a 1,4-dihydropyridine scaffold, only isradipine and benidipine inhibited FATP2, whereas amlodipine and nifedipine were inactive. This divergence indicates that FATP2 inhibition is specific to certain structural features rather than being a general class effect of dihydropyridine calcium channel blockers. Structural comparison reveals that the active compounds are neutral and highly lipophilic, with asymmetric substituents. Conversely, the inactive compounds are either partially charged (amlodipine) or symmetric (nifedipine), suggesting that neutrality, lipophilicity, and asymmetry are critical requirements for binding to site 2.

To further clarify the mechanism of the more potent inhibitor, we determined the structure of FATP2^Benidipine^ at 3.7 Å resolution (Figure 4F; Figure S2). Like isradipine, benidipine traps FATP2 in the thioester-forming conformation and displays a bulky density in site 2. However, additional features in the benidipine density reveal a unique mode of inhibition: the bulky substituent of benidipine extends beyond site 2, protruding into site 1 where it sterically clashes with the primary fatty acid PAM1 (Figure 4G). Consequently, whereas PAM1 is merely repositioned within the tunnel in the FATP2^Isradipine^ structure, the density for PAM1 is entirely absent in the FATP2^Benidipine^complex (Figure 4H). This dual interference mechanism in which benidipine occupies the membrane-proximal Site 2 while simultaneously penetrating the deep Site 1 provides a structural rationale for its superior inhibitory efficacy compared to isradipine. Collectively, these structural insights reveal that isradipine and benidipine function as allosteric inhibitors that physically obstruct the fatty acid binding tunnel. By stabilizing the non-productive thioester-forming conformation and sterically clashing with the substrate at site 1, these agents provide a potent mechanism of inhibition distinct from physiological feedback. This dual mode of action, characterized by site 2 occupation and the enforced displacement or exclusion of the fatty acid, establishes a structural template for the development of targeted FATP2 therapeutics.

### Catalytic cycle and mechanism of vectorial acylation

Integrating our structural snapshots of the adenylate-forming (FATP2^PAM-endo^ and FATP2^ATP–BEM^) and thioester-forming (FATP2^Isradipine^) states allows us to reconstruct the complete catalytic cycle of FATP2 (Figure 5). The reaction trajectory begins with the enzyme in the adenylate-forming conformation in which the N- and C-terminal domains align to create a continuous active site interface. First, a substrate fatty acid enters the secondary fatty acid binding site (site 2) located near the membrane interface adjacent to α2 (Step I). In this state, the hydrophobic tunnel acts as a molecular conduit that translocates the substrate from the site 2 to the catalytic center at the site 1 (Step II) for ATP-dependent adenylation. This results in the formation of the acyl-AMP intermediate (Step III). Following formation of the acyl-AMP intermediate and pyrophosphate release, the enzyme undergoes a substantial structural rearrangement to accommodate coenzyme A. Although we did not directly trap an acyl-AMP–bound state, the domain alternation observed across our adenylate-forming and thioester-forming snapshots is consistent with the canonical ANL reaction trajectory in which acyl-AMP precedes thioesterification. This transition is driven by a ∼130° rotation of the C-terminal domain relative to the N-terminal domain (Step IV). This domain alternation, a hallmark of the ANL superfamily, remodels the active site by exposing the acyl-AMP intermediate to the CoA-binding pocket while shielding the reactive thioester from solvent hydrolysis.^38^ Upon completion of thioesterification, the release of the acyl-CoA product and AMP allows the C-terminal domain to rotate back, resetting the domain architecture to the resting adenylate-forming state (Step V).

**Figure 5.**
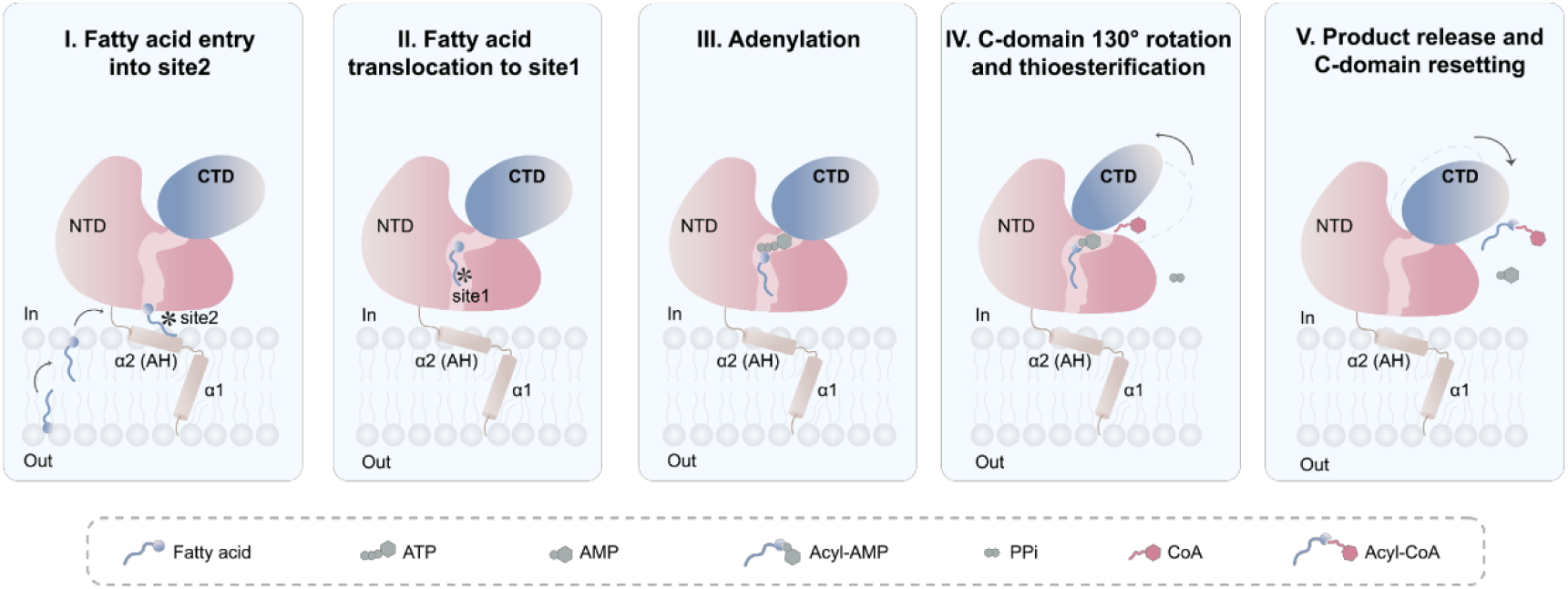
Proposed mechanism of fatty acid uptake and activation by FATP2. Schematic model illustrating the catalytic cycle of FATP2 at the membrane interface. The protein is anchored to the membrane via an N-terminal transmembrane helix (α1) and an amphipathic helix (α2/AH). The catalytic domains (N-terminal domain, NTD; C-terminal domain, CTD) reside in the intracellular space. (I) Entry into site 2: A free fatty acid (blue squiggle) enters the secondary binding pocket (PAM2) from the membrane interface or the intracellular side; (II) Translocation to site 1: The fatty acid translocates through the internal tunnel to the primary active site (site 1); (III) Adenylation: In the presence of ATP (grey circles), the fatty acid is activated to form an acyl-AMP intermediate (Acyl-AMP), releasing pyrophosphate (not shown); (IV) C-domain rotation and thioesterification: The CTD undergoes a large conformational change (approximately 130° rotation) to bring the CoA-binding site (and bound CoA, red squiggle) into proximity with the acyl-AMP intermediate, facilitating the thioesterification reaction; (V) Product release and C-domain resetting: The final acyl-CoA product is released, and AMP is ejected. The CTD rotates back to the open conformation to reset the enzyme for the next cycle.

Collectively, this structural cycle provides the physical basis for the ‘vectorial acylation’ hypothesis. The absence of a canonical transporter fold does not support fatty acid transport via an alternating-access mechanism across the bilayer. Instead, the enzyme functions through vectorial channeling, whereby the hydrophobic tunnel creates a low-energy path for fatty acids to be extracted from the membrane inner leaflet while irreversible catalytic steps driven by ATP hydrolysis provide the thermodynamic pull. By rapidly converting entering fatty acids into membrane-impermeable acyl-CoA, FATP2 maintains a steep concentration gradient that drives continuous uptake independent of a physical translocation pore.

## Discussion

For decades, the identity of FATP2 has been obscured by conflicting evidence, blurring the distinction between a fatty acid transporter and an acyl-CoA synthetase. ^11,20,22,39^ Our structural analysis clarifies this ambiguity: FATP2 is an enzyme that mimics transporter function through a unique membrane-anchored architecture. The "lollipop" topology, tethered by a single transmembrane helix and an amphipathic interface helix, positions the catalytic domain to closely oppose the membrane surface. This contradicts earlier topology models suggesting multi-pass transmembrane domains and aligns FATP2 more closely with the ANL superfamily of enzymes than with the Solute Carrier (SLC) transporters to which it is currently assigned. Consequently, we propose that the "transport" activity attributed to FATP2 is not physical translocation across the bilayer, but rather the result of highly efficient, coupled vectorial acylation that creates a thermodynamic sink to accelerate fatty acid uptake to physiological levels. Central to this mechanism is the identification of a hydrophobic tunnel traversing the N-terminal domain. This feature elucidates how FATP2 overcomes the energetic barrier associated with extracting fatty acids from the membrane bilayer for catalysis in the aqueous cytosol. The tunnel functions as a privileged conduit, sequestering the hydrophobic acyl chain from the solvent environment while guiding the polar carboxyl headgroup toward the ATP-binding site.

This architecture also rationalizes the enzyme’s broad substrate specificity, accommodating bulky pharmacological agents like bempedoic acid and diverse fatty acid chain lengths, provided their acyl tails can navigate the hydrophobic channel.

However, the enzyme is not merely a passive conduit; our study reveals an additional layer of regulation governing this metabolic gatekeeper. The discovery that acyl-CoA products can occupy the ATP-binding pocket and occlude the fatty-acid tunnel represents a direct negative feedback loop, presumably preventing excessive fatty acid accumulation when downstream metabolic demand is low.

Beyond endogenous regulation, our screening reveals that dihydropyridines exploit a unique vulnerability by binding at the interface of the amphipathic helix and the fatty acid tunnel. Crucially, our structural comparison of inhibitor binding reveals a tiered mechanism of blockade. While isradipine effectively displaces the endogenous substrate at the membrane interface (Site 2), benidipine exerts a more profound effect: its bulky scaffold not only occupies Site 2 but extends deep into the protein core, sterically ejecting the fatty acid from the catalytic tunnel (Site 1) entirely. This precise mapping of the inhibitory sites offers a structural foundation for targeting lipid flux in human disease.

Our identification and structural characterization of these inhibitors offer a blueprint for therapeutic intervention. Unlike the endogenous feedback mechanism, these dihydropyridines achieve potent inhibition by locking the enzyme in a non-productive thioester-forming conformation. Given the association of FATP2 with lipid-dependent tumor growth and metabolic diseases, the repurposing of dihydropyridines or the development of isradipine analogs optimized to avoid calcium channel activity may represent a promising strategy to target FATP2-dependent pathologies. In summary, these studies provide a high-resolution view of the machinery coupling fatty acid uptake to activation through the synthesis of acyl-CoAs. By redefining FATP2 as a membrane-anchored enzyme rather than a transporter, we refine current models of cellular lipid acquisition and provide a structural framework for drug discovery.

### Limitations of the study

While our cryo-EM structures successfully resolve the amphipathic helix and transmembrane tether, the use of isolated protein in detergent micelles does not fully recapitulate the complex environment of the cell membrane. Specifically, this experimental setup lacks the native lipid composition and potential protein partners that may cooperate with FATP2 to facilitate fatty acid uptake. Consequently, future investigations employing either lipid-reconstituted systems (e.g., liposomes) or *in situ* structural biology approaches are necessary to determine how the native membrane environment and intermolecular interactions modulate the efficiency of fatty acid extraction via the proposed vectorial channeling mechanism. Second, our structures provide only static snapshots of the adenylate- and thioester-forming steps. As a result, the precise kinetics of the ∼130-degree domain rotation and the dynamic trajectory of the substrate from the membrane interface (site 2) to the catalytic center (site 1) remain to be elucidated. Complementary time-resolved structural methods will be required to capture these transient conformational dynamics.

## Supporting information

Cryo-EM data processing statistics

## ACKNOWLEDGEMENTS

This work was supported by start-up funds from the Guangzhou National Laboratory (Grant NO. GZNL2025C01025 to X.W.), the National Natural Science Foundation of China (Grant No. 32400547 to J.Y. and Grant No. 32501083 to X.W.), the Pearl River Talent Recruitment Program (Grant No. 2023QN10Y296 to J.Y. and Grant No. 2023QN10Y326 to X.W.), and NIH grants R35 GM118090 and R01 CA262055 to R.M. We thank the Guangzhou National Laboratory Core Facility. We thank the staff at the Guangzhou National Laboratory Center for Biological Imaging for their technical support during cryo-EM grid screening and data acquisition. We specifically thank M. Z., M. L., and W. L. for assistance with cryo-EM sample preparation and data collection, and C.Q. for assistance with mass spectrometry analysis.

## AUTHOR CONTRIBUTIONS

Z.Z. and M.Z. purified the FATP2 protein and characterized its enzymatic activity. Z.Z., M.Z. and H.J. collected the cryo-EM data. Z.Z. and M.Z. processed the cryo-EM data and reconstructed the atomic models. Y.H. performed the lipidomic analysis of endogenous ligand bound to FATP2. Z.Z. and X.W. built and refined the structure model. Z.Z. and M.D. performed high-throughput drug screening. Z.Z., M.Z. and X.W. analyzed the structure. X.W., J.S., and R.M. conceived and coordinated the project. Z.Z., M.Z. and X.W. drafted the manuscript, and all authors reviewed, edited, and approved the final version.

## STAR★METHODS

### KEY RESOURCES TABLE

| Reagent or resource | Source | Identifier |
| --- | --- | --- |
| **Cell lines and plasmids** |  |  |
| HEK293F | CRL-3249 | Lab collection |
| pCJ7-*fatp2-flag* | pCJ7 carrying hFATP2-FLAG | This study |
| **Chemical and biological agents** |  |  |
| Anti-DYKDDDDK Affinity Resin | GenScript | L00766-100 |
| n-dodecyl-β-D-maltopyranoside | Anatrace | D310 |
| Cholesteryl hemisuccinate tris salt | Anatrace | CH210 |
| Sodium butyrate | Aladdin | 156-54-7 |
| Butyric acid | MedChemExpress (MCE) | HY-B0350 |
| Decanoic acid | MedChemExpress (MCE) | HY-W015309 |
| Palmitic acid | MedChemExpress (MCE) | HY-N0830 |
| Stearic acid | MedChemExpress (MCE) | HY-B2219 |
| Palmitoleic acid | MedChemExpress (MCE) | HY-W011873 |
| Oleic acid | MedChemExpress (MCE) | HY-N1446 |
| Erucic acid | MedChemExpress (MCE) | HY-N7109 |
| Nervonic acid | MedChemExpress (MCE) | HY-N2526 |
| Bempedoic acid | MedChemExpress (MCE) | HY-12357 |
| ATP | MedChemExpress (MCE) | HY-B2176 |
| Coenzyme A | MedChemExpress (MCE) | HY-128851 |
| 16:0 Coenzyme A triammonium | Merck | 870716P |
| Isradipine | MedChemExpress (MCE) | HY-B0233 |
| Benidipine hydrochloride | MedChemExpress (MCE) | HY-B1448 |
| Amlodipine | MedChemExpress (MCE) | HY-B0317 |
| Nifedipine | MedChemExpress (MCE) | HY-B0284 |
| Bacitracin Zinc | MedChemExpress (MCE) | HY-B0278 |
| Febuxostat | MedChemExpress (MCE) | HY-14268 |
| Aprotinin | MedChemExpress (MCE) | HY-P0017 |
| Leupeptin hemisulfate | MedChemExpress (MCE) | HY-18234A |
| Pepstatin | MedChemExpress (MCE) | HY-P0018 |
| Palmitoleic acid-d<sub>13</sub> | MedChemExpress (MCE) | HY-W011873S1 |
| Oleic acid-<sup>13</sup>C | MedChemExpress (MCE) | HY-N1446S |
| Linoleic acid-<sup>13</sup>C<sub>1</sub> | MedChemExpress (MCE) | HY-N0729S3 |
| 5,5'-dithiobis-(2-nitrobenzoic acid) | Merck | D8130 |

## RESOURCE AVAILABILITY

### Lead contact

Further information and requests for resources and reagents should be directed to, and will be fulfilled by, the lead contact, Xuepeng Wei.

### Materials availability

Plasmids generated in this study are available upon request from the lead contact.

### Data and code availability

- Cryo-EM density maps have been deposited in the Electron Microscopy Data Bank (EMDB) under accession codes EMD-69487 (FATP2^PAM-endo^), EMD-68983 (FATP2^ATP-BEM^), EMD-68974 (FATP2^Product^), EMD-68984 (FATP2^CoA^), EMD-68985 (FATP2^Isradipine^) and EMD-69486 (FATP2^Benidipine^). The corresponding atomic coordinates have been deposited in the Protein Data Bank (PDB) under accession codes 24FY, 23HX, 23HL, 23HY, 23HZ and 24FR respectively.
- This paper does not report original code.
- Any additional information required to reanalyze the data reported in this paper is available from the lead contact upon request.

## EXPERIMENTAL MODEL AND STUDY PARTICIPANT DETAILS

### Cell lines

HEK293F cells were cultured in suspension at 37°C with 8% CO₂. Cells were maintained in FreeStyle 293 Expression Medium and transfected at a density of 2.0–2.5 × 10⁶ cells/mL.

## METHODS DETAILS

### Cloning and protein expression

The cDNA encoding full-length human FATP2 (FATP2; UniProt ID: O14975) was cloned into a modified pCJ7 vector harboring a C-terminal FLAG tag (DYKDDDDK). Wild-type (WT) and mutant constructs generated via site-directed mutagenesis were verified by DNA sequencing (Tsingke Biotechnology). HEK293F cells were cultured in suspension at 37°C with 8% CO_2_ and transfected at a density of 2.0–2.5 × 10⁶ cells/mL. Transfection complexes were generated by incubating 1.3 mg of high-purity plasmid (isolated via Omega Fastfilter Plasmid Maxi Kit) in 50 mL of fresh medium for 5 minutes, followed by the addition of 3 mg linear polyethyleneimine (Polysciences). Following a 15-minute incubation, the mixture was added dropwise to 1 L of cell culture. At 8–12 hours post-transfection, cells were supplemented with 10 mM sodium butyrate (Aladdin) and cultured for an additional 48 hours.

### Protein purification

Cells were harvested by centrifugation (4,000 × g, 5 min) and lysed via sonication in Buffer A (20 mM HEPES pH 7.5, 150 mM NaCl, 2 mM MgCl₂, 5 μg/mL aprotinin (MCE), 2 μg/mL leupeptin hemisulfate (MCE), and 1.4 μg/mL pepstatin (MCE) at 30% power using 10 s on/20 s off cycles. Following lysate clarification via ultracentrifugation (42,000 × g, 45 min, 4°C), the membrane fraction was resuspended in Buffer A supplemented with 1% (w/v) n-dodecyl-β-D-maltopyranoside (DDM; Anatrace) and 0.1% (w/v) cholesteryl hemisuccinate tris salt (CHS; Anatrace). Solubilization was performed for 2 hours at 4°C with continuous agitation. After a second ultracentrifugation step (42,000 × g, 45 min, 4°C), the supernatant was filtered (0.22 μm) and incubated with Anti-DYKDDDDK affinity beads (GenScript) for 1.5 hours at 4°C. The resin was washed with 7 column volumes (CV) of Buffer B (20 mM HEPES pH 7.5, 150 mM NaCl, 2 mM MgCl₂, 0.025% DDM, 0.0025% CHS) and eluted with 6-10 CV of Buffer B supplemented with 0.25 mg/mL 3×FLAG peptide. The eluate was concentrated (100 kDa MWCO) and subjected to size exclusion chromatography (SEC) on a Superose 6 Increase 10/300 GL column equilibrated in Buffer B. Peak fractions were pooled and concentrated for cryo-EM.

### Extraction and LC-MS Analysis of endogenous fatty acids

For lipid extraction, a sample mixture was prepared consisting of 1 mL of protein purification buffer (Buffer B) and 1 mL of FATP2 (2 mg/mL). Internal standards (palmitoleic acid-d13, oleic acid-^13^C, and linoleic acid-^13^C_1_) were added to a final concentration of 60 µM each. The samples were mixed with 2 mL of chloroform-methanol (1:2, v/v) by vortexing, followed by the sequential addition of 0.5 mL of chloroform and 0.5 mL of ultrapure water. The mixture was incubated at room temperature for 2 hours. After incubation, the lipid layer (lower layer) was extracted, dried under a nitrogen stream, and stored at −80°C.

On the day of analysis, the dried lipid extracts were reconstituted in acetonitrile-isopropanol (60:40, v/v). Lipid separation was performed using an Agilent 1290 II ultra-high-performance liquid chromatography (UHPLC) system coupled to a 6546 quadrupole time-of-flight (Q-TOF) mass spectrometer equipped with an electrospray ionization (ESI) source. Chromatography was carried out on an Agilent ZORBAX Eclipse Plus C18 column (2.1 × 50 mm, 1.8 µm) maintained at 45 °C. The mobile phases were: (Mobile Phase A) water-acetonitrile-isopropanol (50:30:20, v/v/v) containing 10 mM ammonium formate and 5 µM methylenediphosphonic acid; and (Mobile Phase B) isopropanol-acetonitrile-water (90:9:1, v/v/v) containing 10 mM ammonium formate. The following gradient program was used: 0 min, 15% Mobile Phase B; 0–1.0 min, increase to 50% Mobile Phase B; 1.0–2.0 min, increase to 100% Mobile Phase B; 2.0–3.0 min, decrease to 15% Mobile Phase B; followed by a 3 min post-run equilibration. The flow rate was 0.40 mL/min and the injection volume was 3 µL.

ESI was operated in negative ion mode with the following settings: nebulizer pressure, 35 psi; sheath gas temperature, 350 °C at 11 L/min; drying gas temperature, 320 °C at 8 L/min; and capillary voltage, 3500 V. The Q-TOF acquisition parameters were: scan range m/z 200–500; scan speed, 1.5 spectra/s; and fragmentor voltage, 175 V. All data were acquired and processed using Agilent MassHunter Workstation software (Data Acquisition version 10.1 and Qualitative Analysis version 10.0; Agilent Technologies, USA).

### FATP2 catalytic activity assay

Catalytic activity was quantified by monitoring the consumption of free Coenzyme A (CoA-SH) using the 5,5′-dithiobis-(2-nitrobenzoic acid) (DTNB) assay. To eliminate background thiol interference, purified FATP2 variants were equilibrated in a non-reducing buffer composed of 20 mM HEPES (pH 7.5), 150 mM NaCl, 2 mM MgCl₂, 0.025% (w/v) DDM, and 0.0025% (w/v) CHS. Enzymatic reactions were performed in a final volume of 25 µL containing 1 µM FATP2 protein, 100 µM CoA, 1 mM bempedoic acid, 100 µM ATP, and 0.1 mg/mL BSA, and incubated at room temperature for 1 hour. Subsequently, a 16 µL aliquot of the reaction mixture was quenched by the addition of 24 µL of detection reagent, which consisted of 2 mM DTNB dissolved in a solution of 50 mM sodium acetate and 1 M Tris (pH 8.0) mixed at a 1:2 (v/v) ratio. After a 5 min incubation, absorbance was measured at 412 nm using a BioTek spectrophotometer. Catalytic activity was calculated based on the reduction in free thiols relative to a substrate-only negative control, with wild-type FATP2 activity normalized to 100%, and data are presented as mean ± s.d. (*n* ≥3 technical replicates).

### High-throughput screening for FATP2 inhibitors

The catalytic assay was adapted for high-throughput screening (HTS) using a library of FDA-approved drugs. Screening was performed in 384-well microplates to monitor the acyl-CoA synthetase activity of FATP2. Briefly, 0.26 µL of each compound (5 mM stock) was dispensed into wells containing 12.5 µL of recombinant FATP2 (2 µM) via pin-tool, resulting in a final compound concentration of 50 µM. Reactions were initiated by the addition of 12.5 µL of a substrate mixture containing ATP, CoA, and bempedoic acid. The final concentrations in the 25 µL reaction volume were 1 µM FATP2 and 500 µM for each substrate (ATP, CoA, and bempedoic acid). The mixture was incubated at room temperature for 1 hour.

Controls were included in each assay plate: positive controls contained the complete reaction mixture, while negative controls excluded ATP to establish the baseline signal. Following incubation, 37.5 µL of DTNB detection reagent was added directly to each well. After 5 min, absorbance was measured at 412 nm using a microplate reader.

### Hit validation and IC_50_ determination

Assay robustness was demonstrated by a Z′-factor of 0.7, indicating a high-quality screening window. Primary hits were identified based on significant inhibition of CoA consumption, manifested as retention of absorbance signal relative to the active controls. Following manual curation to exclude potential false positives and pan-assay interference compounds (PAINS), 12 candidates advanced to the secondary screening. Confirmatory assays performed in triplicate identified four compounds with reproducible inhibitory activity. Subsequently, these four compounds were selected for dose–response analysis to determine their half-maximal inhibitory concentrations (IC_50_).

### Cryo-EM sample preparation and data acquisition

Purified FATP2 (5–10 mg/mL) containing endogenous fatty acid was prepared for vitrification either in the absence of exogenous ligands or following a 1-hour incubation on ice with specific ligands, including: (1) 0.5 mM ATP supplemented with 0.2 mM bempedoic acid; (2) 1 mM AMP and 0.5 mM 16:0 Coenzyme A triammonium (Merck); (3) 0.5 mM Coenzyme A; or (4) 0.25 mM Isradipine (MCE); (5) 0.2 mM Benidipine (MCE). Aliquots (2.5 µL) were applied to glow-discharged UltrAuFoil Au 300 1.2/1.3 grids, blotted for 3.5 s at 100% humidity, and vitrified in liquid ethane using a Vitrobot Mark IV. Data acquisition was performed on two 300 kV Titan Krios microscopes, both equipped with Falcon 4i direct electron detectors. Imaging was conducted at a nominal magnification of 215,000×, corresponding to calibrated pixel sizes of 0.572 Å and 0.578 Å, respectively. Dose-fractionated movies (40 frames; total accumulated dose of 50 e⁻/Å²) were collected using EPU software with a defocus range of −0.8 to −1.6 μm.

### Cryo-EM data processing

Data processing was performed using CryoSPARC^40,41^. Beam-induced motion was corrected using Patch Motion Correction, and gain correction was applied using the facility-supplied reference. Contrast transfer function (CTF) parameters were estimated via Patch CTF; micrographs exhibiting a CTF fit resolution worse than 6 Å or astigmatism greater than 500 Å were excluded from the dataset. The overall data processing workflow is presented in Figure S2.

For the FATP2^PAM-endo^ complex, initial particle selection was performed on a representative subset of 904 micrographs using the Blob Picker. These particles were subjected to reference-free 2D classification to remove bad particles, followed by ab initio reconstruction and heterogeneous refinement to generate an initial 3D reference. Particles from the optimal class were subjected to Non-Uniform Refinement to yield a consensus map, from which 50 equally spaced projections were generated. These projections were employed as templates to pick particles from the full dataset of 7,683 micrographs, resulting in the extraction of 2,013,736 particles with a box size of 480 pixels (Fourier-cropped by a factor of 4 for initial processing). To efficiently clean the dataset, particles were partitioned into three subsets and subjected to multiple rounds of seed-facilitated heterogeneous refinement using the initial consensus map as a reference. The best particles were recombined and re-extracted at the full box size of 480 pixels (unbinned), yielding a consensus map at 3.4 Å resolution comprising 38,659 particles. Visual inspection of the consensus map revealed a homodimeric architecture; however, the reconstruction exhibited marked asymmetry in map quality. While one protomer was structurally ordered with well-defined features, the neighboring protomer displayed attenuated density consistent with significant continuous conformational heterogeneity. Attempts to refine the dataset with C2 symmetry imposed resulted in a degradation of map quality and lower resolution. This outcome indicates that the structural plasticity of the disordered protomer, or stochastic variations at the dimer interface, breaks the strict non-crystallographic symmetry required for averaging.

Consequently, to isolate the high-resolution signal, we performed local refinement using a mask encompassing only the ordered protomer. By excluding the incoherent signal contributed by the flexible neighbor, we improved the alignment accuracy, resulting in a final reconstruction at 3.2 Å resolution.

The FATP2^ATP-BEM^, FATP2^Product,^ and FATP2^CoA^ datasets were processed using an analogous workflow, yielding global consensus maps at 2.6 Å, 2.7 Å, and 3.3 Å resolution, respectively.

For the FATP2^Isradipine^ and FATP2^Benidipine^ datasets, the consensus maps displayed diffuse and poorly resolved density in the C-terminal region, indicating conformational heterogeneity distinct from the other datasets. To resolve this, we performed 3D classification without alignment using a mask covering the FATP2 protein (excluding the detergent micelle). This strategy successfully separated the particles into two distinct conformational populations: an adenylate-forming state and a unique thioester-forming state. To improve the map quality and resolution of these distinct conformers, particles from each class were subjected to local refinement. For the FATP2^Isradipine^ dataset, this resulted in final reconstructions at 4.6 Å (adenylate-forming) and 3.5 Å (thioester-forming) resolution. Similarly, the FATP2^Benidipine^ dataset yielded final reconstructions at 4.4 Å (adenylate-forming) and 3.7 Å (thioester-forming) resolution. The 4.6 Å and 4.4 Å reconstructions exhibit an adenylate-forming conformation identical to that observed in the FATP2^PAM-endo^ and FATP2^ATP-BEM^ structures; however, due to their limited resolution compared to the other datasets, no atomic models were built for these specific maps.

### Model building and refinement

For model building and refinement, the N- and C-terminal domains of the AlphaFold-predicted human FATP2 structure were initially docked into the cryo-EM density maps using ChimeraX^42^. These models were subsequently adjusted manually in COOT using the real-space refinement module^43^. The coordinates were subjected to iterative rounds of real-space refinement in PHENIX^44^. Model quality was validated via PHENIX, assessing clash scores, MolProbity scores, and Ramachandran statistics. Structural statistics are summarized in Table 1. Figures were prepared using ChimeraX or PyMOL.

## QUANTIFICATION AND STATISTICAL ANALYSIS

Statistical analyses were performed using GraphPad Prism 10. Quantification methods and tools used are described in each relevant section of the methods or figure legends.

## Supplementary Figures

**Figure S1.**
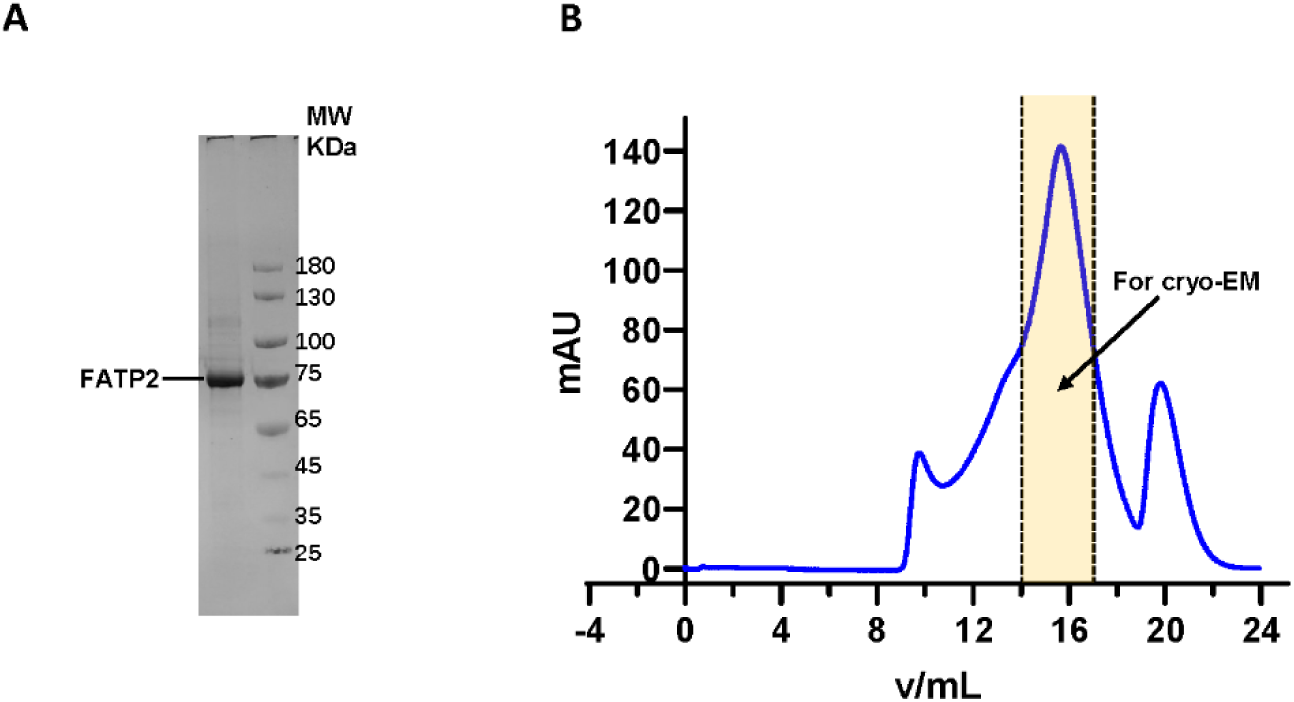
Purification and characterization of FATP2, related to Figure 1. **(A)** SDS-PAGE analysis of purified FATP2. The lane corresponds to the peak fraction eluted at ∼16 mL from size exclusion chromatography (SEC), which was concentrated for cryo-EM analysis. Molecular weight markers (kDa) are indicated on the right. **(B)** Representative SEC profile of FATP2 on a Superose 6 Increase 10/300 GL column, showing the elution peak at approximately 16 mL.

**Figure S2.**
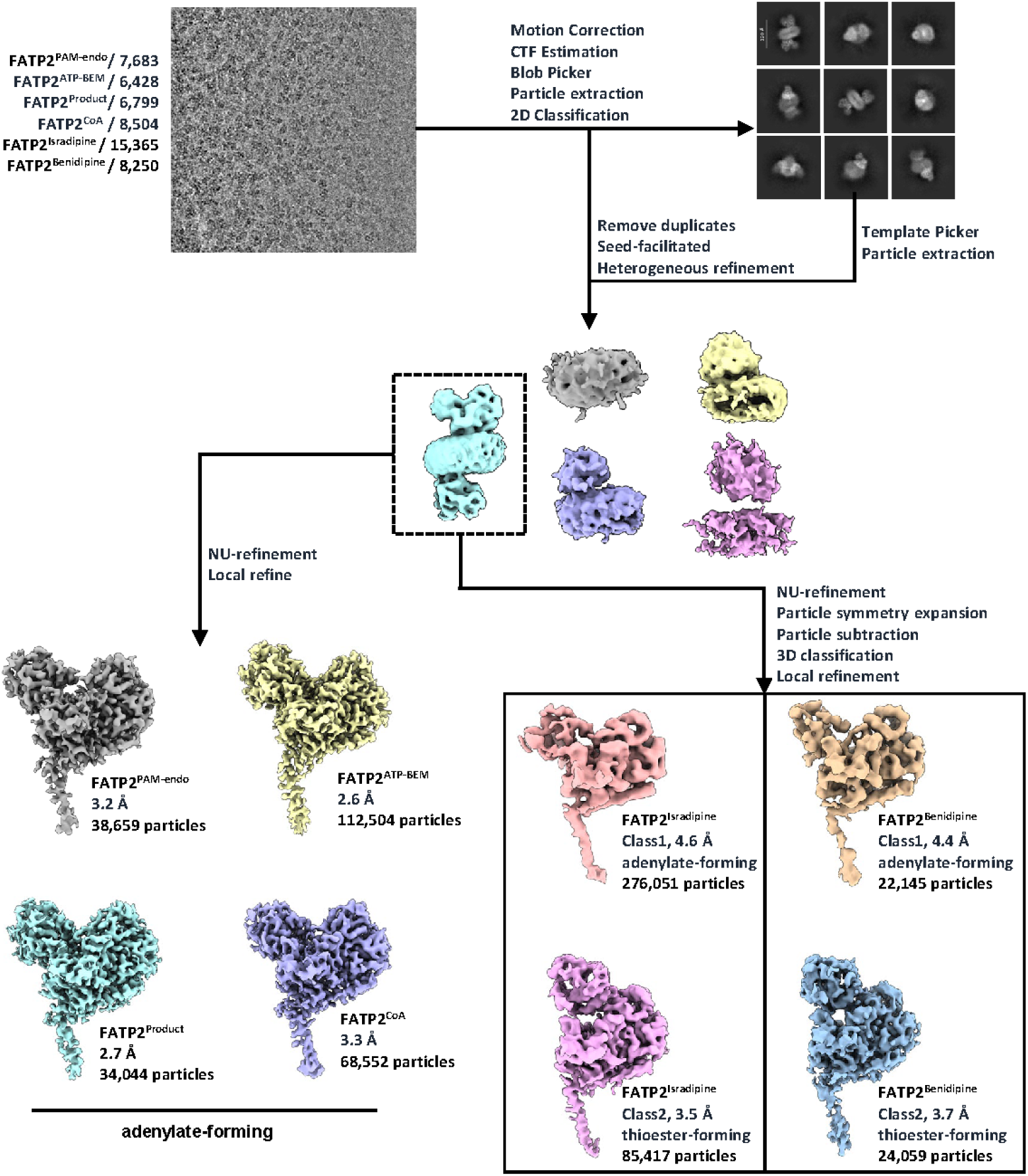
Workflow of FATP2 cryo-EM data processing, related to Figures 1-4.

**Figure S3.**
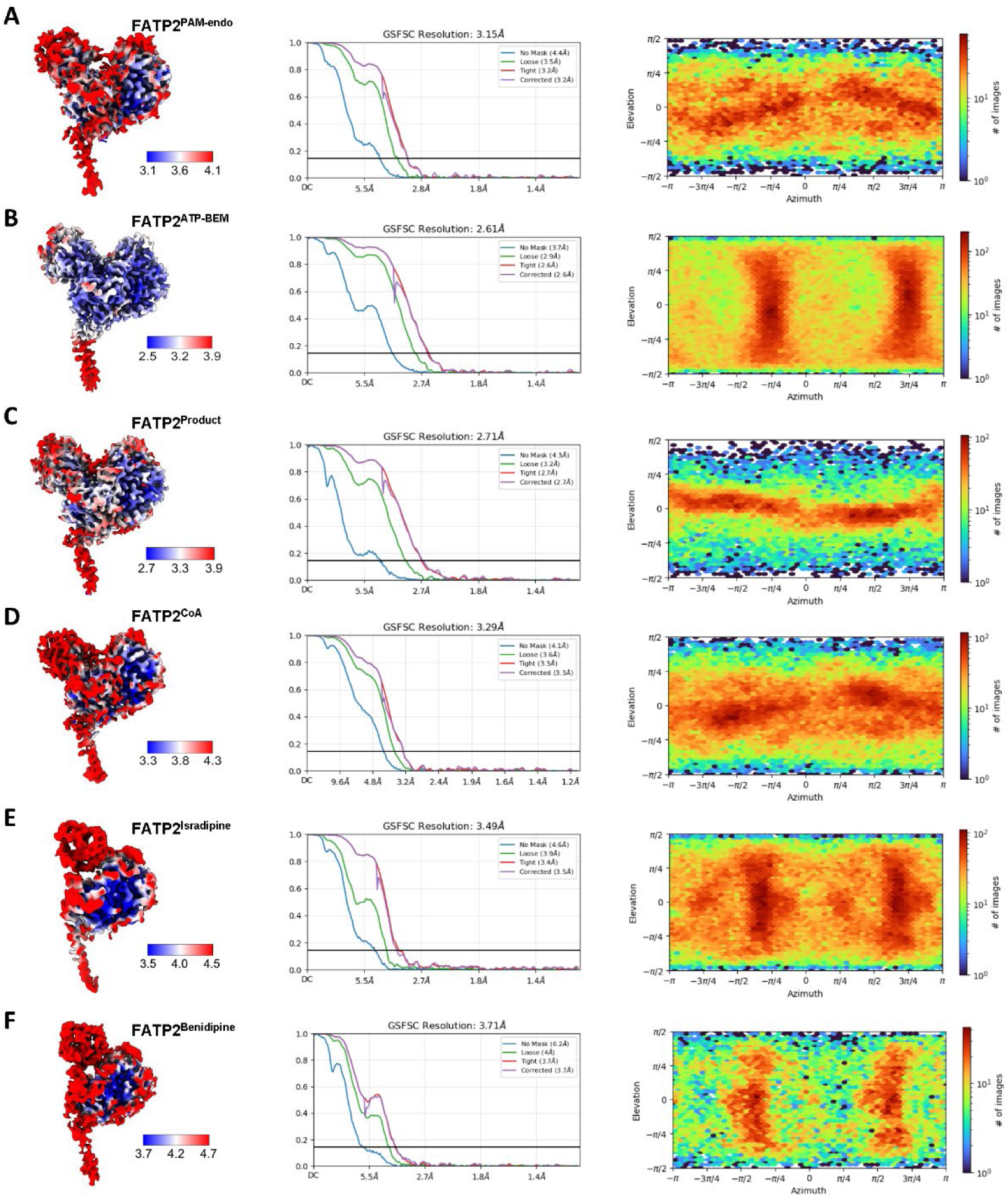
Cryo-EM resolution estimation and validation for FATP2 complexes, related to Figures 1-4. (A to F) Cryo-EM analysis of FATP2^PAM-endo^ (A), FATP2^ATP-BEM^ (B), FATP2^Product^ (C), FATP2^CoA^ (D), FATP2^Isradipine^ (E), and FATP2^Benidipine^ (F). For each state, the panels display: (Left) Local resolution map colored according to the resolution scale bar (in Å), where blue indicates higher resolution and red indicates lower resolution. (Middle) Gold-standard Fourier shell correlation (FSC) curves. The global resolution is determined at the FSC = 0.143 criterion (indicated by the horizontal black line). (Right) Angular distribution of the particles used for the final reconstruction, showing the orientation coverage.

**Figure S4.**
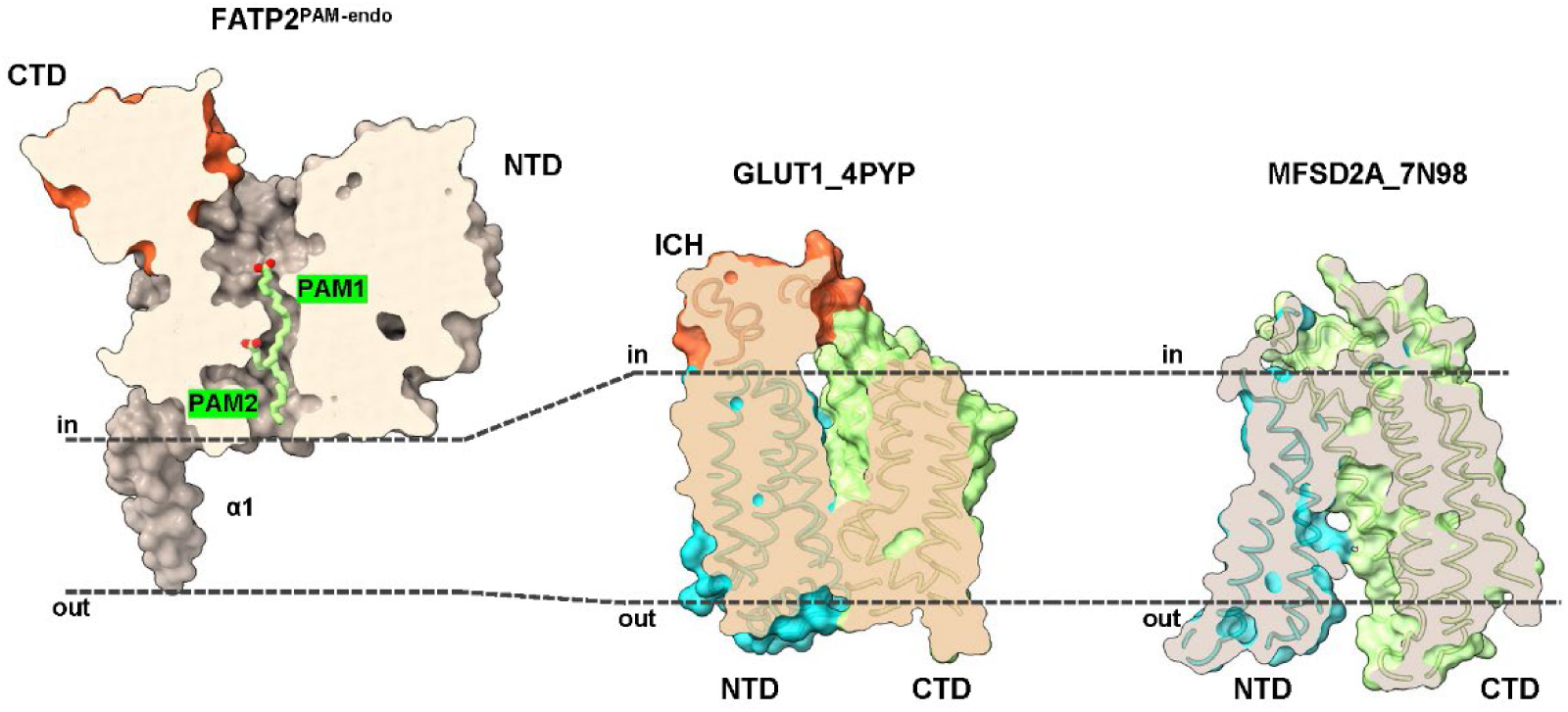
Structural divergence of FATP2 from canonical transporters, related to Figure 1. Comparison with GLUT1 (SLC2A1) and MFSD2A highlights that FATP2 lacks the typical transmembrane architecture defined by established glucose and lipid transporters.

**Figure S5.**
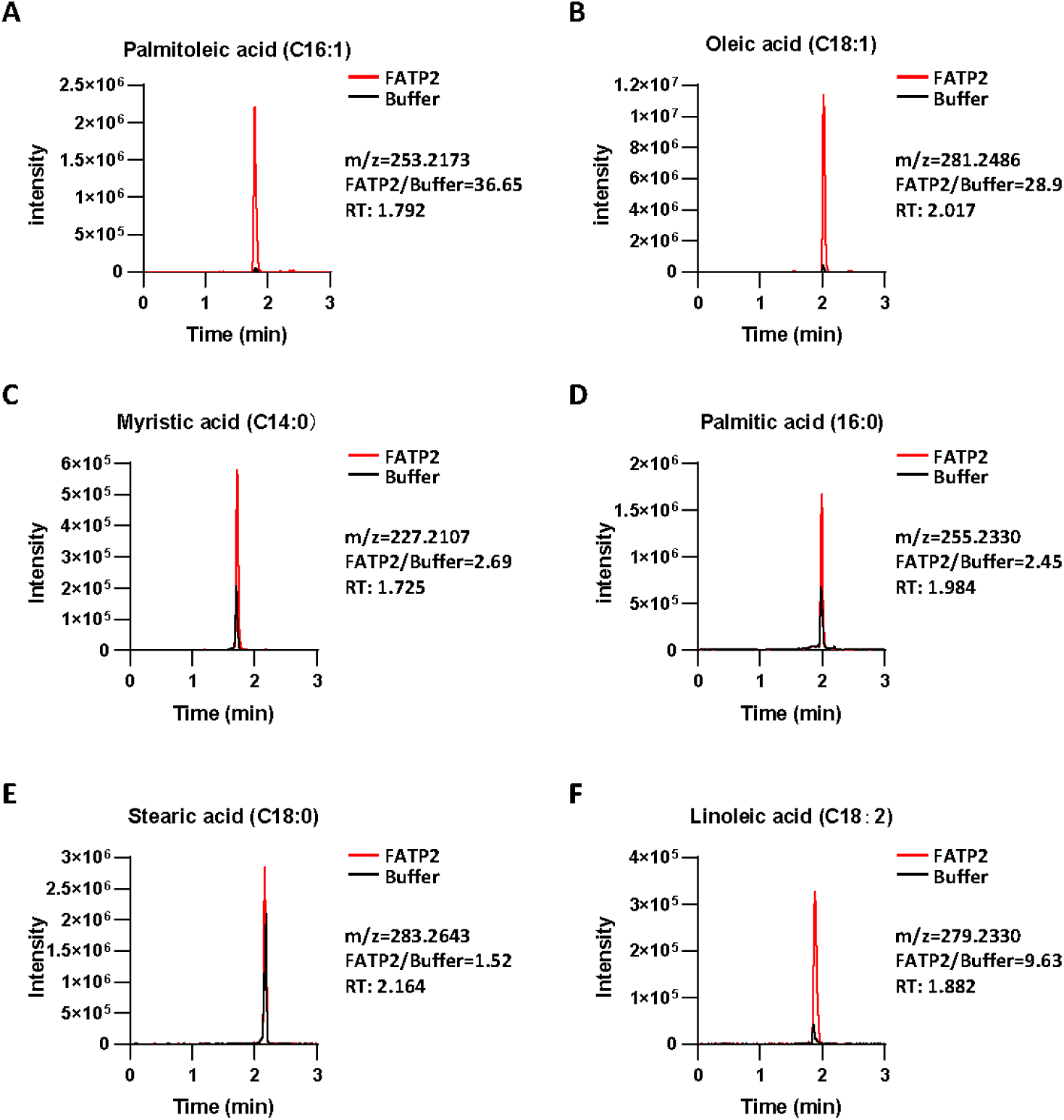
LC-MS analysis of endogenous fatty acids copurified with FATP2, related to Figure 1. (A–F) Extracted ion chromatograms comparing purified FATP2 (red) versus buffer control (black) for: (A) Palmitoleic acid (C16:1); (B) Oleic acid (C18:1); (C) Myristic acid (C14:0); (D) Palmitic acid (C16:0); (E) Stearic acid (C18:0); and (F) Linoleic acid (C18:2). Insets provide the mass-to-charge ratio (m/z), the fold enrichment ratio of signal intensity in the FATP2 sample versus buffer (FATP2/Buffer), and the retention time (RT) for each peak. The data indicate a strong preference for mono-unsaturated fatty acids (Palmitoleic and Oleic acid) binding to FATP2 compared to saturated fatty acids.

**Figure S6.**
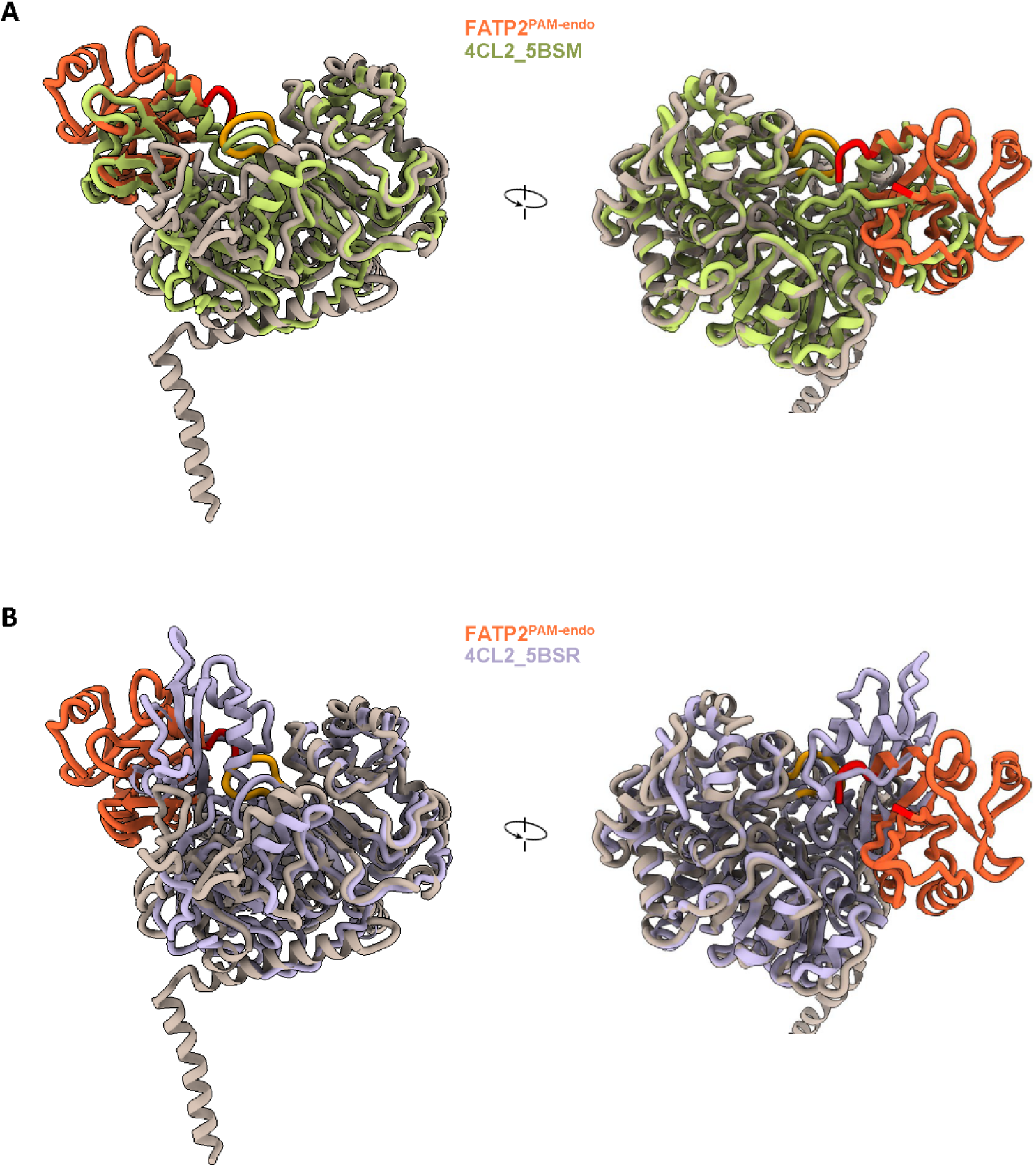
Structural comparison of FATP2^PAM-endo^ with plant 4-coumarate:CoA ligase 2 (4CL2), related to Figure 1. (A) Superposition of FATP2^PAM-endo^ (orange) with the adenylate-forming state of 4CL2 (PDB: 5BSM, lime green). The C-terminal domain of FATP2 aligns closely with that of 4CL2 in the adenylate-forming conformation. (B) Superposition of FATP2^PAM-endo^ (orange) with the thioester-forming state of 4CL2 (PDB: 5BSR, light purple). The significant deviation of the C-terminal domain indicates that FATP2 does not adopt the thioester-forming conformation in this structure.

**Figure S7.**
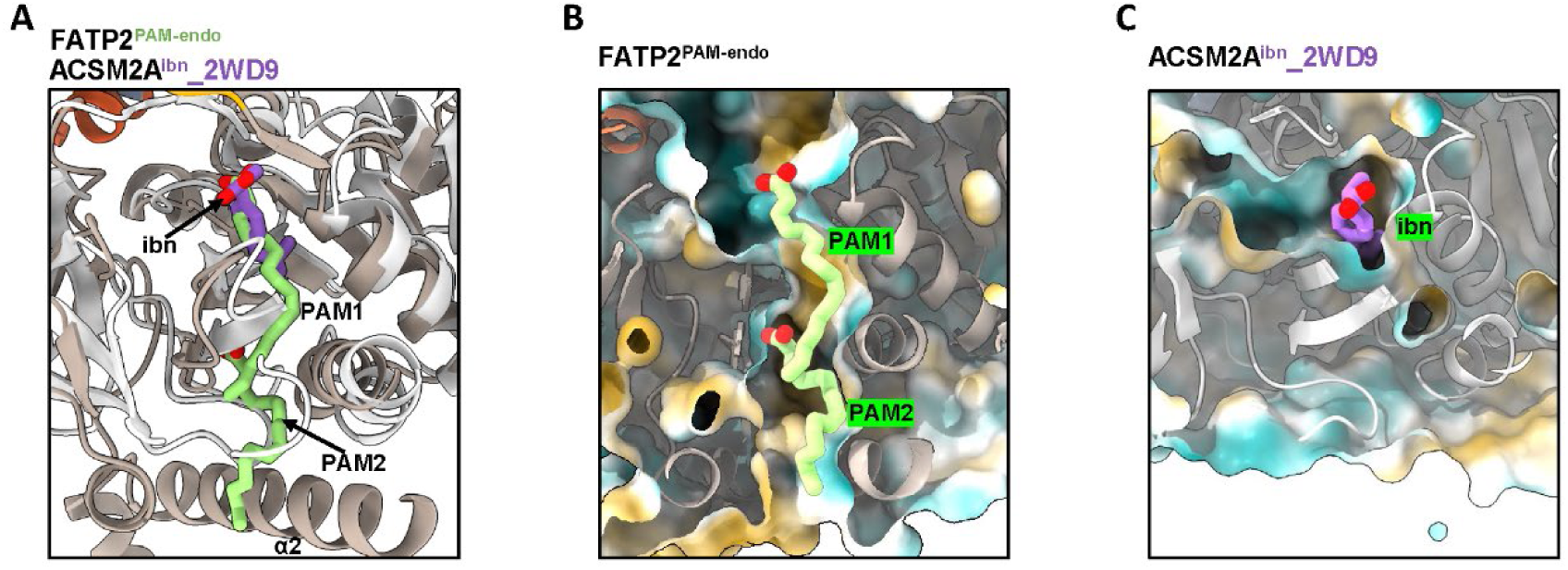
Comparison of FATP2 and ACSM2A substrate-binding sites, related to Figure 1. (A) Superposition of FATP2^PAM-endo^ (grey) and ACSM2A (brown; PDB ID: 2WD9). Ligands are shown as sticks: endogenous fatty acids (PAM1/PAM2) in FATP2 are green; the ibuprofen-AMP analog (ibn) in ACSM2A is purple. (B) Cut-away surface of the FATP2 tunnel colored by hydrophobicity (gold, hydrophobic; cyan, hydrophilic). The long, continuous tunnel accommodates two fatty acids (PAM1 and PAM2). (C) Cut-away surface of the ACSM2A pocket (PDB ID: 2WD9), colored as in (B). The ACSM2A pocket is shallower and obstructed, consistent with specificity for medium-chain or bulky substrates.

**Figure S8.**
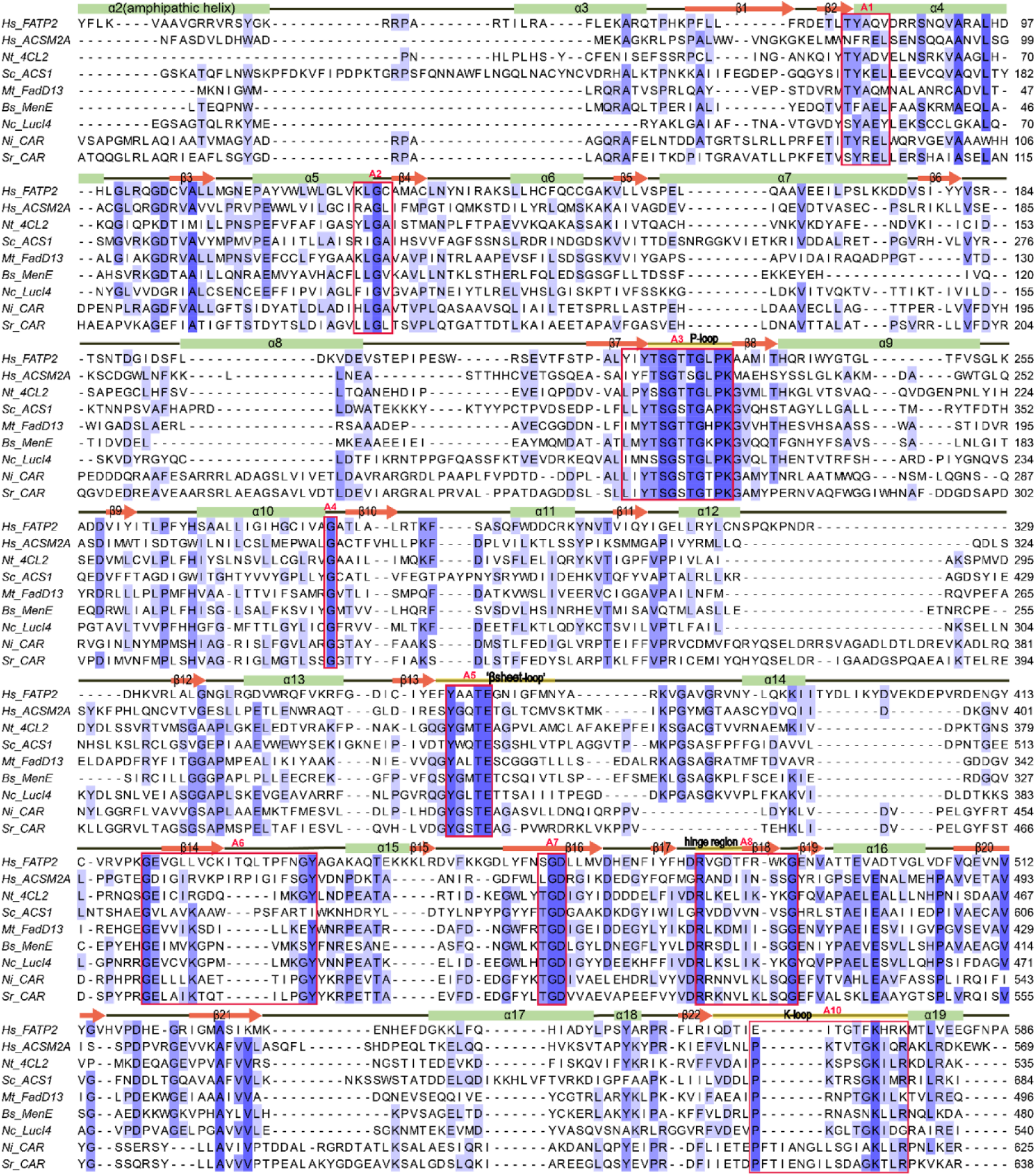
Multiple sequence alignment of *Homo sapiens* FATP2 with representative ANL superfamily members, related to Figure 2. Secondary structure elements corresponding to the human FATP2 structure are depicted above the alignment (spirals, α-helices; arrows, β-strands). Residues are shaded based on conservation, with dark blue indicating identity and light blue indicating similarity. Conserved core motifs (A1–A10) characteristic of the ANL superfamily are outlined in red boxes. Notably, the phosphate-binding P-loop (motif A3) and the essential K-loop (motif A10), which contain key residues strictly required for nucleotide binding and coordination, exhibit a high degree of conservation across all aligned sequences.

**Figure S9.**
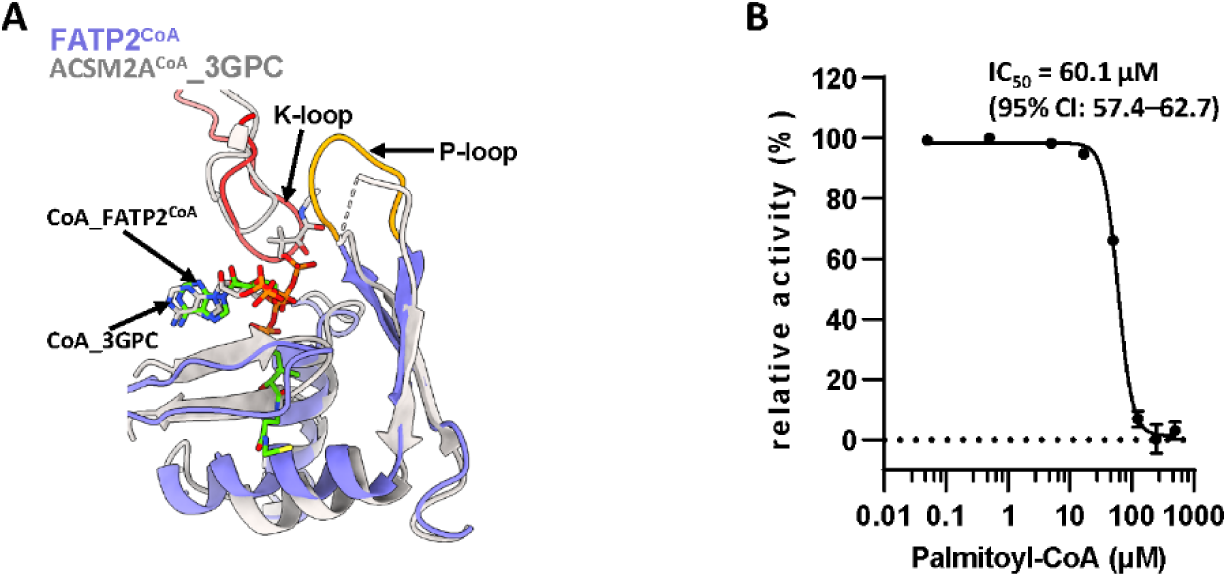
Structural comparison of FATP2^CoA^ with ACSM2A-CoA and feedback inhibition of FATP2 by palmitoyl-CoA, related to Figure 3. (A) Structural superposition of the FATP2^CoA^ complex (slate blue ribbons) and the ACSM2A–CoA complex (grey ribbons; PDB ID: 3GPC). The phosphate-binding loop (P-loop) is colored yellow, and the loop containing the catalytic lysine (K-loop) is colored red. The CoA molecules are shown in stick representation, highlighting the distinct conformation of the cofactor in FATP2^CoA^, green carbons) compared to the bent conformation observed in ACSM2A (CoA_3GPC, grey carbons). (B) Dose-response curve showing the inhibition of FATP2 enzymatic activity by palmitoyl-CoA. Relative activity is plotted against increasing concentrations of palmitoyl-CoA (log scale). The calculated half-maximal inhibitory concentration (IC_50_) is 60.1 µM (95% confidence interval: 57.4–62.7 µM).

**Figure S10.**
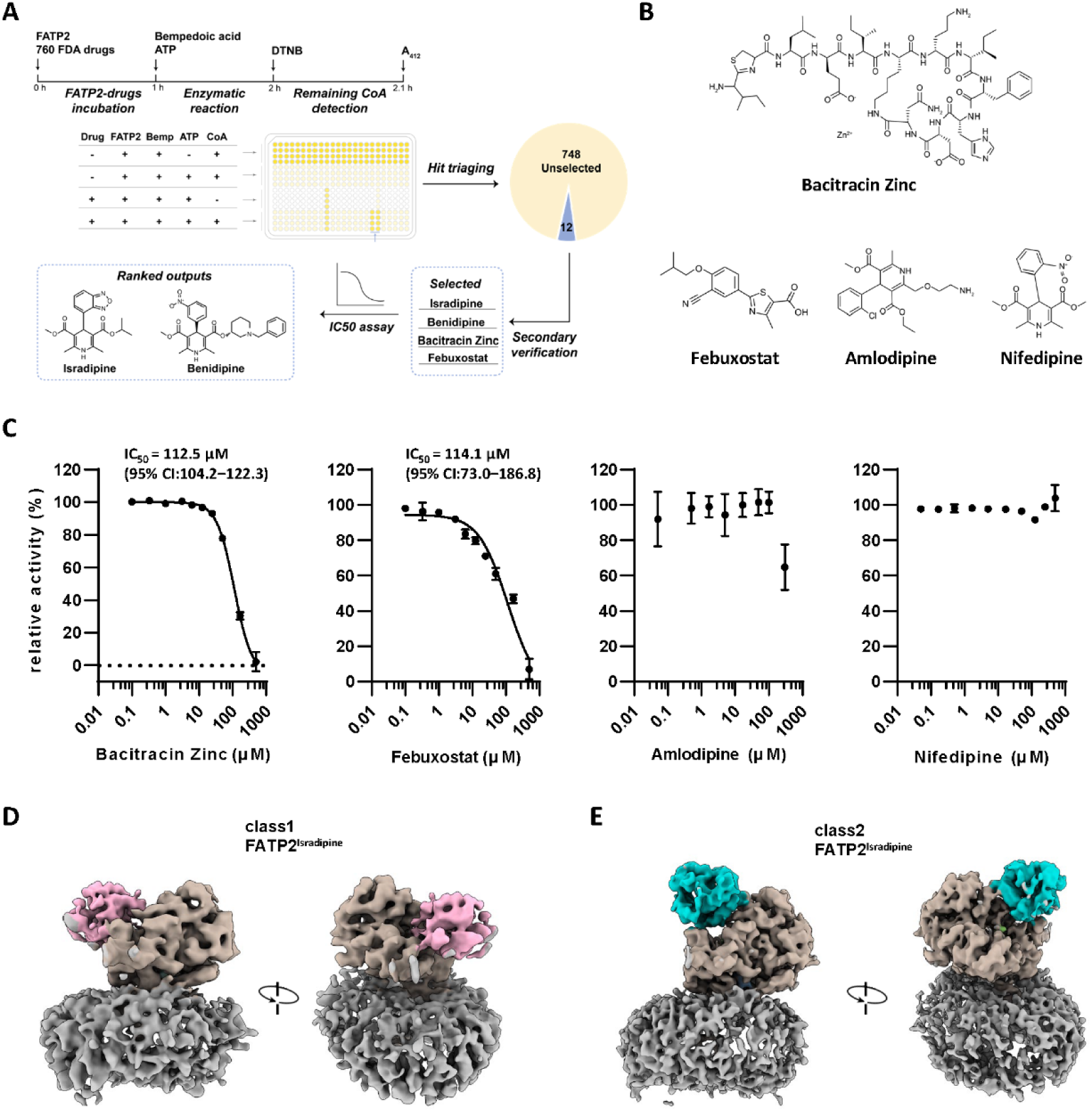
Identification of FATP2 inhibitors via high-throughput screening and structural characterization, related to Figure 4. (A) Schematic workflow of the high-throughput screening (HTS) campaign. A library of 760 FDA-approved drugs was screened for FATP2 inhibition by monitoring residual Coenzyme A (CoA) levels using a DTNB (Ellman’s reagent) colorimetric assay at 412 nm. Of the 760 compounds, 12 primary hits were identified and subjected to secondary verification and IC_50_ determination to select final candidates. (B) Chemical structures of the identified hits Bacitracin Zinc and Febuxostat, alongside the structural analogs Amlodipine and Nifedipine included for specificity analysis. (C) Dose–response analysis of low-potency hits and structural analogs. (Left two panels) Bacitracin Zinc and Febuxostat were identified in the primary screen but displayed low potency, with IC_50_ values of 112.5 µM (95% CI: 104.2–122.3) and 114.1 µM (95% CI: 73.0–186.8), respectively. Due to this weak inhibition compared to Isradipine and Benidipine, they were excluded from further structural characterization. (Right two panels) Amlodipine and Nifedipine were subsequently tested due to their structural similarity to Isradipine (dihydropyridine analogs). Despite this similarity, they displayed no inhibitory activity, demonstrating the strict structural specificity required for binding. Data are presented as mean ± s.d. (n ≥3 technical replicates). (D and E) Cryo-EM 3D classification of the FATP2^Isradipine^ complex. The analysis reveals conformational heterogeneity with two distinct classes: class 1 (D), showing the flexible domain in a distinct orientation (pink), and class 2 (E), showing an alternative orientation (cyan). Orthogonal views are shown for each class.

**Figure S11.**
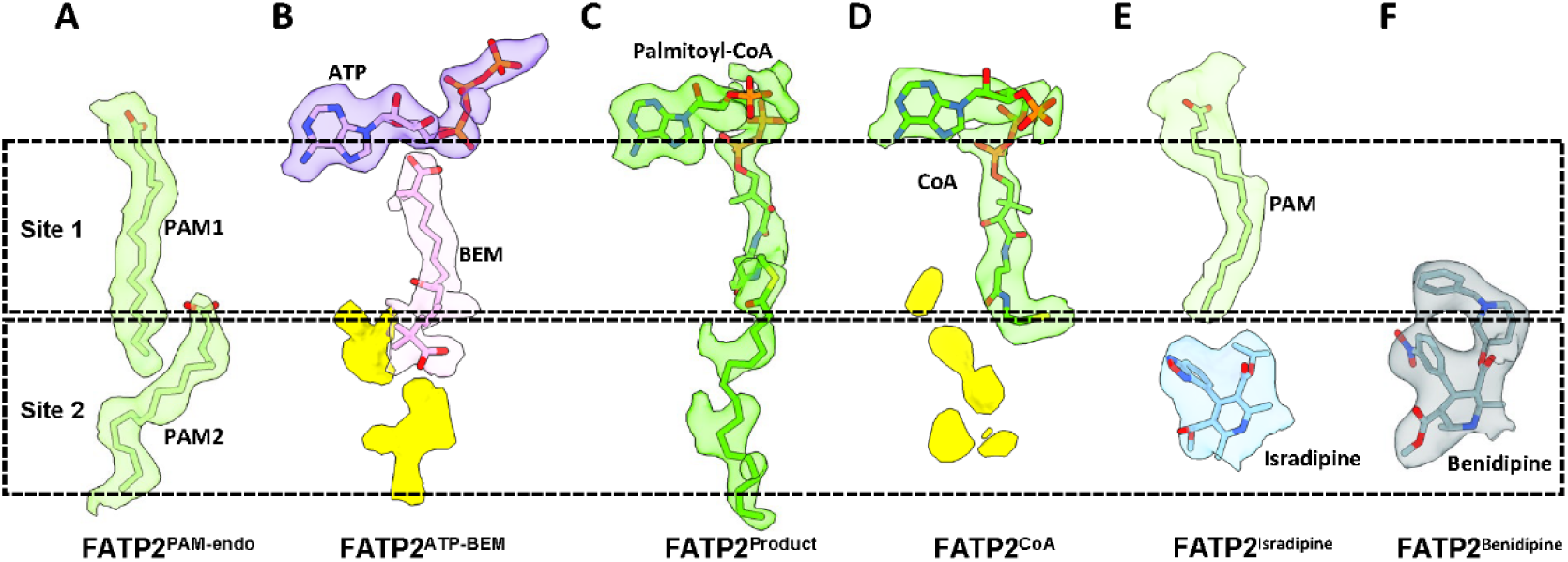
Cryo-EM density of ligands occupying Site 1 and Site 2 in FATP2 structures, related to Figures 1-5. Cryo-EM density maps (transparent surface) and atomic models (sticks) for ligands bound to FATP2. Horizontal dashed lines demarcate the boundaries of the primary substrate tunnel (Site 1, corresponding to the PAM1 site) and the secondary binding pocket (Site 2, corresponding to the PAM2 site). (A) FATP2^PAM-endo^: density corresponding to endogenous fatty acids (modeled as palmitoleic acid, PAM) occupying both Site 1 and Site 2. (B) FATP2^ATP–BEM^: density for ATP and bempedoic acid. The adenosine moiety is highlighted with a purple transparent surface, and the bempedoic acid moiety with a pink transparent surface. Unmodeled density features in Site 2 are colored yellow. (C) FATP2^Product^: continuous density for the product palmitoyl-CoA, which spans the entire length of the binding channel from the ATP-binding region through Site 1 and into Site 2. (D) FATP2^CoA^: density for Coenzyme A (CoA) bound in the upper active site region. Discontinuous yellow surfaces in Site 2 indicate unmodeled density. (E) FATP2^Isradipine^: density showing an endogenous fatty acid occupying Site 1, while the drug isradipine (highlighted with a light blue transparent surface) is specifically bound in Site 2. (F) FATP2^Benidipine^: density for the drug benidipine (highlighted with a grey transparent surface) bound in Site 2.

## REFERENCES

1. Nakamura, M.T., Yudell, B.E., and Loor, J.J. (2014). Regulation of energy metabolism by long-chain fatty acids. Prog Lipid Res 53, 124–144. 10.1016/j.plipres.2013.12.001.

2. Wei, Y., Wang, D., Topczewski, F., and Pagliassotti, M.J. (2006). Saturated fatty acids induce endoplasmic reticulum stress and apoptosis independently of ceramide in liver cells. Am J Physiol Endocrinol Metab 291, E275–281. 10.1152/ajpendo.00644.2005.

3. Steinberg, S.J., Morgenthaler, J., Heinzer, A.K., Smith, K.D., and Watkins, P.A. (2000). Very long-chain acyl-CoA synthetases. Human "bubblegum" represents a new family of proteins capable of activating very long-chain fatty acids. J Biol Chem 275, 35162–35169. 10.1074/jbc.M006403200.

4. Zou, Z., Tong, F., Faergeman, N.J., Borsting, C., Black, P.N., and DiRusso, C.C. (2003). Vectorial acylation in Saccharomyces cerevisiae. Fat1p and fatty acyl-CoA synthetase are interacting components of a fatty acid import complex. J Biol Chem 278, 16414–16422. 10.1074/jbc.M210557200.

5. Sozio, M.S., Liangpunsakul, S., and Crabb, D. (2010). The role of lipid metabolism in the pathogenesis of alcoholic and nonalcoholic hepatic steatosis. Semin Liver Dis 30, 378–390. 10.1055/s-0030-1267538.

6. Hoy, A.J., Nagarajan, S.R., and Butler, L.M. (2021). Tumour fatty acid metabolism in the context of therapy resistance and obesity. Nat Rev Cancer 21, 753–766. 10.1038/s41568-021-00388-4.

7. Martin-Perez, M., Urdiroz-Urricelqui, U., Bigas, C., and Benitah, S.A. (2022). The role of lipids in cancer progression and metastasis. Cell Metab 34, 1675–1699. 10.1016/j.cmet.2022.09.023.

8. Veglia, F., Tyurin, V.A., Blasi, M., De Leo, A., Kossenkov, A.V., Donthireddy, L., To, T.K.J., Schug, Z., Basu, S., Wang, F., et al. (2019). Fatty acid transport protein 2 reprograms neutrophils in cancer. Nature 569, 73–78. 10.1038/s41586-019-1118-2.

9. Stahl, A. (2004). A current review of fatty acid transport proteins (SLC27). Pflugers Arch 447, 722–727. 10.1007/s00424-003-1106-z.

10. Falcon, A., Doege, H., Fluitt, A., Tsang, B., Watson, N., Kay, M.A., and Stahl, A. (2010). FATP2 is a hepatic fatty acid transporter and peroxisomal very long-chain acyl-CoA synthetase. Am J Physiol Endocrinol Metab 299, E384–393. 10.1152/ajpendo.00226.2010.

11. Schaap, F.G., Hamers, L., Van der Vusse, G.J., and Glatz, J.F. (1997). Molecular cloning of fatty acid-transport protein cDNA from rat. Biochim Biophys Acta 1354, 29–34. 10.1016/s0167-4781(97)00121-8.

12. Gimeno, R.E. (2007). Fatty acid transport proteins. Curr Opin Lipidol 18, 271–276. 10.1097/MOL.0b013e3281338558.

13. Nissen, S.E., Lincoff, A.M., Brennan, D., Ray, K.K., Mason, D., Kastelein, J.J.P., Thompson, P.D., Libby, P., Cho, L., Plutzky, J., et al. (2023). Bempedoic Acid and Cardiovascular Outcomes in Statin-Intolerant Patients. N Engl J Med 388, 1353–1364. 10.1056/NEJMoa2215024.

14. Pinkosky, S.L., Newton, R.S., Day, E.A., Ford, R.J., Lhotak, S., Austin, R.C., Birch, C.M., Smith, B.K., Filippov, S., Groot, P.H.E., et al. (2016). Liver-specific ATP-citrate lyase inhibition by bempedoic acid decreases LDL-C and attenuates atherosclerosis. Nat Commun 7, 13457. 10.1038/ncomms13457.

15. Gautam, J., Wu, J., Lally, J.S.V., McNicol, J.D., Fayyazi, R., Ahmadi, E., Oniciu, D.C., Heaton, S., Newton, R.S., Rehal, S., et al. (2025). ACLY inhibition promotes tumour immunity and suppresses liver cancer. Nature 645, 507–517. 10.1038/s41586-025-09297-0.

16. Black, P.N., and DiRusso, C.C. (2025). FATP2 at the crossroads of fatty acid transport, lipotoxicity, and complex disease. J Clin Invest 135. 10.1172/JCI199873.

17. Anderson, C.M., and Stahl, A. (2013). SLC27 fatty acid transport proteins. Mol Aspects Med 34, 516–528. 10.1016/j.mam.2012.07.010.

18. Lewis, S.E., Listenberger, L.L., Ory, D.S., and Schaffer, J.E. (2001). Membrane topology of the murine fatty acid transport protein 1. J Biol Chem 276, 37042–37050. 10.1074/jbc.M105556200.

19. Obermeyer, T., Fraisl, P., DiRusso, C.C., and Black, P.N. (2007). Topology of the yeast fatty acid transport protein Fat1p: mechanistic implications for functional domains on the cytosolic surface of the plasma membrane. J Lipid Res 48, 2354–2364. 10.1194/jlr.M700300-JLR200.

20. Schaffer, J.E., and Lodish, H.F. (1994). Expression cloning and characterization of a novel adipocyte long chain fatty acid transport protein. Cell 79, 427–436. 10.1016/0092-8674(94)90252-6.

21. Jumper, J., Evans, R., Pritzel, A., Green, T., Figurnov, M., Ronneberger, O., Tunyasuvunakool, K., Bates, R., Zidek, A., Potapenko, A., et al. (2021). Highly accurate protein structure prediction with AlphaFold. Nature 596, 583–589. 10.1038/s41586-021-03819-2.

22. Jia, Z., Pei, Z., Maiguel, D., Toomer, C.J., and Watkins, P.A. (2007). The fatty acid transport protein (FATP) family: very long chain acyl-CoA synthetases or solute carriers? J Mol Neurosci 33, 25–31. 10.1007/s12031-007-0038-z.

23. Fitton, A., and Benfield, P. (1990). Isradipine. A review of its pharmacodynamic and pharmacokinetic properties, and therapeutic use in cardiovascular disease. Drugs 40, 31–74. 10.2165/00003495-199040010-00004.

24. Yamamoto, E., Kataoka, K., Dong, Y.F., Nakamura, T., Fukuda, M., Nako, H., Ogawa, H., and Kim-Mitsuyama, S. (2010). Benidipine, a dihydropyridine L-type/T-type calcium channel blocker, affords additive benefits for prevention of cardiorenal injury in hypertensive rats. J Hypertens 28, 1321–1329. 10.1097/HJH.0b013e3283388045.

25. Schmelz, S., and Naismith, J.H. (2009). Adenylate-forming enzymes. Curr Opin Struct Biol 19, 666–671. 10.1016/j.sbi.2009.09.004.

26. Colas, C., Ung, P.M., and Schlessinger, A. (2016). SLC Transporters: Structure, Function, and Drug Discovery. Medchemcomm 7, 1069–1081. 10.1039/C6MD00005C.

27. Deng, D., Sun, P., Yan, C., Ke, M., Jiang, X., Xiong, L., Ren, W., Hirata, K., Yamamoto, M., Fan, S., and Yan, N. (2015). Molecular basis of ligand recognition and transport by glucose transporters. Nature 526, 391–396. 10.1038/nature14655.

28. Schlessinger, A., Khuri, N., Giacomini, K.M., and Sali, A. (2013). Molecular modeling and ligand docking for solute carrier (SLC) transporters. Curr Top Med Chem 13, 843–856. 10.2174/1568026611313070007.

29. Deng, D., Xu, C., Sun, P., Wu, J., Yan, C., Hu, M., and Yan, N. (2014). Crystal structure of the human glucose transporter GLUT1. Nature 510, 121–125. 10.1038/nature13306.

30. Wood, C.A.P., Zhang, J., Aydin, D., Xu, Y., Andreone, B.J., Langen, U.H., Dror, R.O., Gu, C., and Feng, L. (2021). Structure and mechanism of blood-brain-barrier lipid transporter MFSD2A. Nature 596, 444–448. 10.1038/s41586-021-03782-y.

31. Li, Z., and Nair, S.K. (2015). Structural Basis for Specificity and Flexibility in a Plant 4-Coumarate:CoA Ligase. Structure 23, 2032–2042. 10.1016/j.str.2015.08.012.

32. Kochan, G., Pilka, E.S., von Delft, F., Oppermann, U., and Yue, W.W. (2009). Structural snapshots for the conformation-dependent catalysis by human medium-chain acyl-coenzyme A synthetase ACSM2A. J Mol Biol 388, 997–1008. 10.1016/j.jmb.2009.03.064.

33. Jezewski, A.J., Esan, T.E., Propp, J., Fuller, A.J., Daraji, D.G., Lail, C., 3rd, Staker, B.L., Woodward, E.L., Liu, L., Battaile, K.P., et al. (2024). A single Leishmania adenylate-forming enzyme of the ANL superfamily generates both acetyl- and acetoacetyl-CoA. J Biol Chem 300, 107879. 10.1016/j.jbc.2024.107879.

34. Hall, A.M., Smith, A.J., and Bernlohr, D.A. (2003). Characterization of the Acyl-CoA synthetase activity of purified murine fatty acid transport protein 1. J Biol Chem 278, 43008–43013. 10.1074/jbc.M306575200.

35. Grossman, E., Messerli, F.H., Oren, S., Nunez, B., and Garavaglia, G.E. (1991). Cardiovascular effects of isradipine in essential hypertension. Am J Cardiol 68, 65–70. 10.1016/0002-9149(91)90712-t.

36. Perry, H.M., Jr., Hall, W.D., Benz, J.R., Bartels, D.W., Kostis, J.B., Townsend, R.R., Due, D.L., Peng, A., and Sirgo, M. (1994). Efficacy and safety of atenolol, enalapril, and isradipine in elderly hypertensive women. Am J Med 96, 77–86. 10.1016/0002-9343(94)90118-x.

37. Miyashita, Y., Peterson, D., Rees, J.M., and Flynn, J.T. (2010). Isradipine for treatment of acute hypertension in hospitalized children and adolescents. J Clin Hypertens (Greenwich) 12, 850–855. 10.1111/j.1751-7176.2010.00347.x.

38. Gulick, A.M. (2009). Conformational dynamics in the Acyl-CoA synthetases, adenylation domains of non-ribosomal peptide synthetases, and firefly luciferase. ACS Chem Biol 4, 811–827. 10.1021/cb900156h.

39. Black, P.N., Ahowesso, C., Montefusco, D., Saini, N., and DiRusso, C.C. (2016). Fatty Acid Transport Proteins: Targeting FATP2 as a Gatekeeper Involved in the Transport of Exogenous Fatty Acids. Medchemcomm 7, 612–622. 10.1039/C6MD00043F.

40. Punjani, A., Rubinstein, J.L., Fleet, D.J., and Brubaker, M.A. (2017). cryoSPARC: algorithms for rapid unsupervised cryo-EM structure determination. Nat Methods 14, 290–296. 10.1038/nmeth.4169.

41. Punjani, A., Zhang, H., and Fleet, D.J. (2020). Non-uniform refinement: adaptive regularization improves single-particle cryo-EM reconstruction. Nat Methods 17, 1214–1221. 10.1038/s41592-020-00990-8.

42. Pettersen, E.F., Goddard, T.D., Huang, C.C., Meng, E.C., Couch, G.S., Croll, T.I., Morris, J.H., and Ferrin, T.E. (2021). UCSF ChimeraX: Structure visualization for researchers, educators, and developers. Protein Sci 30, 70–82. 10.1002/pro.3943.

43. Emsley, P., and Cowtan, K. (2004). Coot: model-building tools for molecular graphics. Acta Crystallogr D Biol Crystallogr 60, 2126–2132. 10.1107/S0907444904019158.

44. Adams, P.D., Afonine, P.V., Bunkoczi, G., Chen, V.B., Davis, I.W., Echols, N., Headd, J.J., Hung, L.W., Kapral, G.J., Grosse-Kunstleve, R.W., et al. (2010). PHENIX: a comprehensive Python-based system for macromolecular structure solution. Acta Crystallogr D Biol Crystallogr 66, 213–221. 10.1107/S0907444909052925.

