## Supplementary material for "Mechanism of fatty acid uptake and inhibition in human FATP2": Cryo-EM data processing statistics

**Table S1**

**Cryo-EM data collection, processing, and refinement statistics, related to Figures 1-4**

| **Structure** | FATP2$^{PAM\text{-}endo}$ | FATP2$^{ATP+BEM}$ | FATP2$^{Product}$ | FATP2$^{CoA}$ | FATP2$^{Isradipine}$ | FATP2$^{benidipine}$ |
|---|---|---|---|---|---|---|
| **EMDB** | EMD-69487 | EMD-68983 | EMD-68974 | EMD-68984 | EMD-68985 | EMD-69486 |
| **PDB** | 24FY | 23HX | 23HL | 23HY | 23HZ | 24FR |
| **Data collection and processing** | | | | | | |
| Magnification | 215,000 | 215,000 | 215,000 | 215,000 | 215,000 | 215,000 |
| Voltage (kV) | 300 | 300 | 300 | 300 | 300 | 300 |
| Electron exposure (e–/Å$^2$) | 50 | 50 | 50 | 50 | 50 | 50 |
| Defocus range (μm) | -0.8 ~ -1.6 | -0.8 ~ -1.6 | -0.8 ~ -1.6 | -0.8 ~ -1.6 | -0.8 ~ -1.6 | -0.8 ~ -1.6 |
| Pixel size (Å) | 0.578 | 0.572 | 0.572 | 0.578 | 0.572 | 0.572 |
| Symmetry imposed | C1 | C1 | C1 | C1 | C1 | C1 |
| Initial particle images (no.) | 2,013,736 | 3,061,270 | 2,170,235 | 5,676,537 | 8,593,619 | 4,268,809 |
| Final particle images (no.) | 38,659 | 112,504 | 34,044 | 68,552 | 85,417 | 24,059 |
| Map resolution (Å) | 3.2 | 2.6 | 2.7 | 3.3 | 3.5 | 3.7 |
| FSC threshold | 0.143 | 0.143 | 0.143 | 0.143 | 0.143 | 0.143 |
| **Refinement** | | | | | | |
| Initial model used (PDB code) | AlphaFold2 | FATP2$^{PAM\text{-}endo}$ | FATP2$^{PAM\text{-}endo}$ | FATP2$^{PAM\text{-}endo}$ | FATP2$^{PAM\text{-}endo}$ | FATP2$^{PAM\text{-}endo}$ |
| Model resolution (Å) | 3.3 | 2.7 | 2.9 | 3.4 | 3.7 | 3.9 |
| FSC threshold | 0.5 | 0.5 | 0.5 | 0.5 | 0.5 | 0.5 |
| Map sharpening *B* factor (Å$^2$) | -83.7 | -63.8 | -54.1 | -92.8 | -77.8 | -81.5 |
| Model composition | | | | | | |
| Non-hydrogen atoms | 4,925 | 5,013 | 5,023 | 5,006 | 5,003 | 4,995 |
| Protein residues | 611 | 620 | 620 | 620 | 620 | 620 |
| Ligands | 2 | 2 | 1 | 1 | 2 | 1 |
| *B* factors (Å$^2$) | | | | | | |
| Protein | 68.43 | 44.07 | 49.62 | 88.53 | 197.02 | 115.58 |
| Ligand | 62.72 | 47.01 | 57.47 | 86.27 | 139.09 | 76.42 |
| R.m.s. deviations | | | | | | |
| Bond lengths (Å) | 0.003 | 0.002 | 0.006 | 0.003 | 0.003 | 0.004 |
| Bond angles (°) | 0.471 | 0.472 | 0.999 | 0.554 | 0.674 | 0.753 |
| Validation | | | | | | |
| MolProbity score | 2.11 | 1.78 | 2.06 | 2.51 | 2.33 | 2.02 |
| Clashscore | 7.77 | 7.19 | 7.75 | 11.17 | 14.77 | 14.93 |
| Poor rotamers (%) | 3.64 | 2.64 | 3.21 | 3.77 | 2.92 | 0.57 |
| Ramachandran plot | | | | | | |
| Favored (%) | 96.21 | 97.73 | 96.28 | 91.26 | 94.82 | 95.15 |
| Allowed (%) | 3.62 | 2.27 | 3.56 | 8.74 | 4.85 | 4.69 |
| Disallowed (%) | 0.16 | 0.00 | 0.16 | 0.00 | 0.32 | 0.16 |
